# Active transport of tRNAs facilitates distributed protein synthesis

**DOI:** 10.64898/2026.01.26.698744

**Authors:** Jennifer M. Petrosino, Vasiliki Courelli, Keita Uchida, Charles Bond, Barry Cooperman, Alexey Bogush, Melike Lakadamyali, Benjamin L. Prosser

## Abstract

Large, polarized cells such as cardiomyocytes, skeletal myofibers, and neurons rely on localized protein synthesis to sustain size, remodel and adapt to stress. The subcellular architecture of these cells is also inherently unfavorable for long-range, diffusion-based transport, which may promote their heavy reliance on active transport mechanisms for the localization of RNA and proteins. Transfer RNAs (tRNAs) function as essential regulators of protein synthesis by linking transcription and translation. Since their discovery in the 1950s, tRNA subcellular distribution has been assumed to occur through passive diffusion. Here, we report that there are pools of tRNAs that depend on the microtubule network for distribution in cardiomyocytes, skeletal myofibers and neurons. Employing dual-color live and fixed-cell super-resolution imaging, we demonstrate that active transport of tRNAs involves hitchhiking on the exterior of endolysosomal vesicles (ELV). We establish that leucyl-tRNA synthetase (LeuRS), the tRNA-binding protein that charges leucine to its cognate tRNA and interacts with Rag GTP on the surface of ELVs, is essential for tRNA transport. Disruption of LeuRS-ELV interactions is sufficient to impair long-range, microtubule-dependent tRNA transport, without affecting mRNA or rRNA transport. We also show that preventing tRNA transport is sufficient to impair translation at sites distal from the nucleus as well as globally impair protein synthesis, ultimately reducing cell size. These findings redefine tRNAs as actively trafficked cargo and establish their directed transport as a fundamental layer of translation regulation required for myocyte homeostasis.

**Highlights:** 1. Pools of tRNAs require an intact microtubule network for distribution in large, complex cells.

2. tRNAs undergo active transport by hitchhiking on endolysosomal vesicles trafficked by microtubule motors.

3. The tRNA synthetase LeuRS functions as a tRNA-specific adaptor linking tRNAs to endolysosomal vesicles, and disrupting this interaction selectively impairs tRNA, but not mRNA or rRNA transport.

4. Loss of tRNA transport reduces distal translation, global protein synthesis, and cell size.

## Introduction

Large, highly structured cells like striated muscle cells and neurons share a common challenge: their unique subcellular environments are dominated by rigid and highly polarized architecture that is inherently unfavorable for long-range, diffusion-based transport. As terminally differentiated cells, when they remodel in response to physiological and pathological cues, muscle cells change in shape and size, and neurons change by reshaping their synaptic and axonal domains. This process of changing muscle and neuronal shape and size depends on robust and spatially controlled protein synthesis.

It has long been established that mRNAs maintain distinct subcellular localizations as a means to control gene expression^1^ (reviewed in^2^). mRNAs and translational machinery undergo active transport, enabling compartmentalized translation in specialized cellular domains. While active mRNA transport has been extensively studied in neurons (reviewed in ^2,3^), recent work from our group^4^ and others^5^ have shown that striated muscle cells also actively transport and locally translate mRNAs to achieve spatiotemporal control of protein synthesis. mRNA transport is generally thought to occur through ribonucleoprotein (RNP) granules that move along microtubules via motor proteins, although recent evidence indicates that some mRNAs can also hitchhike on the surface of endolysosomal vesicles (ELVs)^6^. Notably, RNPs associated with motile endosomes support protein synthesis at sites far from the neuronal soma, suggesting that RNA transport and distributed translation are closely intertwined.

By contrast, the spatial organization of transfer RNAs (tRNAs) remains almost completely unexplored. tRNAs are small non-coding RNAs that link transcription to translation by reading mRNAs’ codons and transferring cognate amino acids. Since the 1950s^7^, tRNAs are assumed to be distributed by rapid diffusion-based mechanisms. This assumption is largely derived from single molecule imaging studies in live prokaryotic and eukaryotic cells showing that tRNAs exhibit predominantly rapid, stochastic diffusion, or transient slow states during translation-dependent ribosome-binding events^8–12^. Thus, the current dogma is that tRNAs do not require dedicated transport machinery and that they function mainly as available substrates waiting to participate in protein synthesis.

Here, we show that there are pools of tRNAs that undergo active transport in mammalian cells by hitchhiking on endolysosomal (ELV) vesicles, a pathway operant in cardiomyocytes, skeletal myofibers, and neurons. We found that microtubule-motor dependent transport of tRNAs requires the tRNA binding protein LeuRS for the association and co-transport of these RNAs with ELVs. Disruptions to the microtubule network, Kinesin-1 or LeuRS, led to impaired tRNA localization, reduced hitchhiking events, disrupted translation, and myocyte atrophy. Specifically disrupting the interaction between LeuRS and ELVs disrupts microtubule-dependent transport of tRNAs without effecting mRNA or rRNA trafficking. Our findings here redefine tRNAs as actively trafficked cargo that utilize a unique transport mechanism to support local protein synthesis and cellular homeostasis.

## Results

### tRNAs require microtubules for their localization in terminally differentiated cells

We first sought to design and validate tools to monitor tRNA subcellular localization in mammalian cells. We generated hairpin chain reaction (HCR) fluorescent in situ hybridization (FISH) probes to detect either single tRNA species (Gly^GCC^, Gly^CCC^, Pro^AGG^) or the top 50 most abundant tRNAs in striated muscle^13^ (pan tRNA probe) using a pooled probe set (**Figures 1A and S1A**). The tRNA probes exhibited discrete puncta that were non-randomly distributed throughout the cardiomyocyte or skeletal myofiber. As tRNAs are transcribed by Polymerase III (Pol3) and have a half-life of ∼96h^14^, we validated probe specificity by treating adult cardiomyocytes for 96 h with 10 µM of a Pol3 inhibitor, which largely eliminated any signal from either the specific or pan tRNA probes, while polyA mRNA levels were unperturbed (**Figures 1B, S1B, and S1C**).

**Figure 1.**
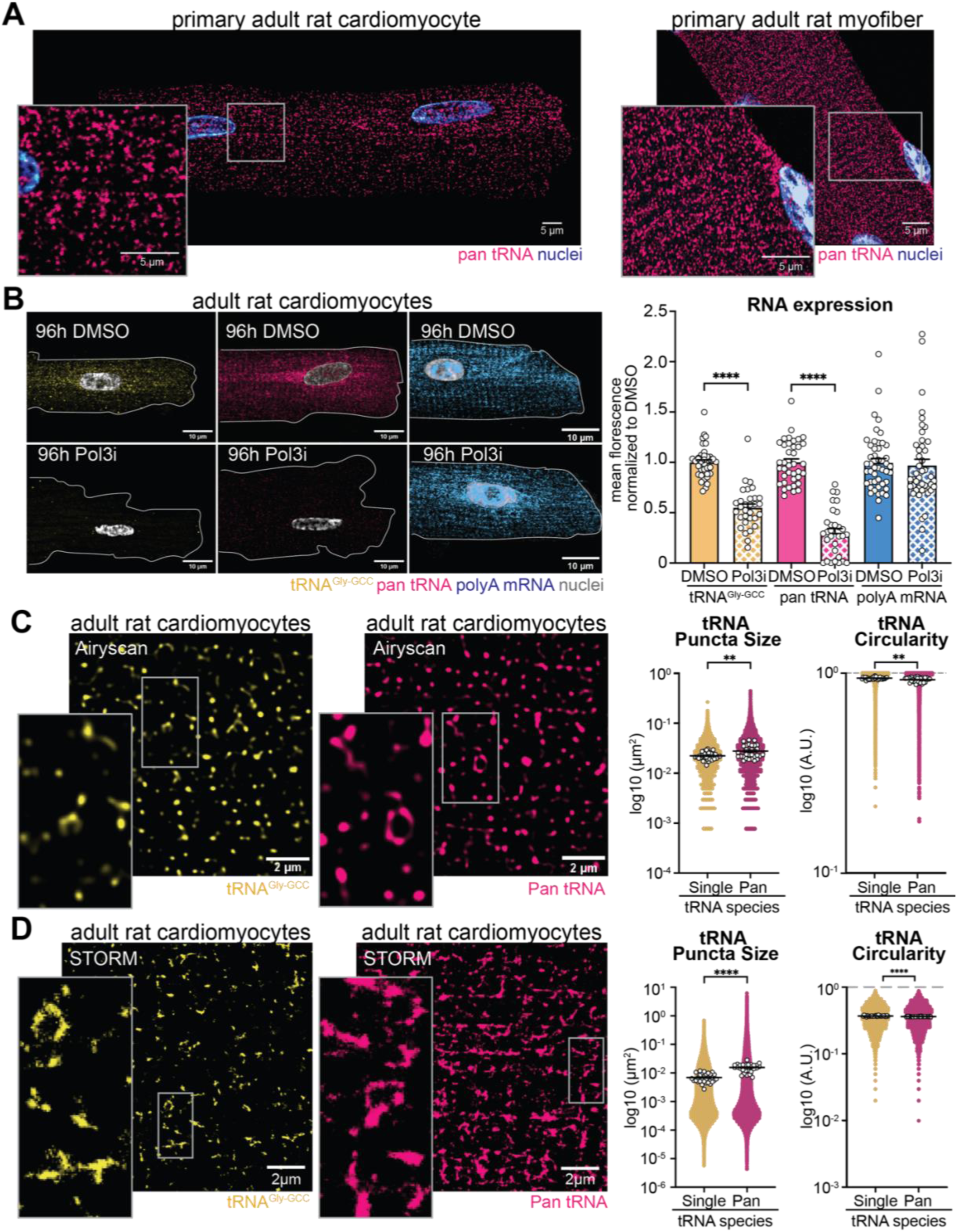
Visualization of endogenous tRNAs in mammalian cells. (A) Representative images of pan tRNA probe signal (pink) and nuclear signal (Hoechst, blue) in primary adult rat ventricular cardiomyocyte (ARVM) and primary rat flexor digitorum brevis (FDB) myofiber. (B) Representative images of tRNA^Gly-GCC^ (yellow), pan tRNA (pink), and polyA mRNA (blue) probes in ARVMs treated for 96h with DMSO or a Polymerase III inhibitor (10 μM Pol3i) and quantification of cellular mean fluorescence intensity normalized to DMSO. (C) Representative high-resolution Airyscan with deconvolution and (D) super-resolution STORM images of tRNA^Gly-GCC^ (yellow) and pan tRNA (pink), and quantification of tRNA puncta size and circularity plotted on a log10 scale, with each individual graphical dot indicating an individual puncta measurement and white dots indicating means for each individual cell. All experiments were done in biological triplicate (*N* = 3), with 10 cells imaged per biological replicate (*n* = 10). Data are presented as the mean ± SEM. *P < 0.05, **P < 0.01, ***P < 0.001, ****P < 0.0001. White boxes indicate magnified fields of view from respective images. Two-sided Student’s t-test was used in Pol3i experiments and against the mean averages of tRNA size and shape in Airyscan and STORM experiments.

We next performed high resolution Airyscan microscopy and quantitative analysis of single and pan tRNA probe signals (**Figure 1C**). tRNAs exhibited a range of unanticipated sizes and shapes, the latter measured by circularity, where 1 indicates a perfect circle and decreasing values indicate a more elongated shape. tRNA puncta often displayed elongated, tubular, or occasionally vesicular morphologies (**Figure 1C**). Puncta detected using the pan tRNA probe were notably larger and more elongated than those detected by the single tRNA probe, suggesting multi-tRNA containing complexes (**Figure 1C**). To orthogonally validate these observations and probe these morphological features at higher resolution, we utilized stochastic optical reconstruction microscopy (STORM) nanoscale imaging (**Figure 1D**). Larger, more elongated morphologies were more apparent using non-diffraction limited microscopy, where clusters of discrete tRNA puncta formed complex assemblies (**Figure 1D**). Nanoscale imaging validated that clusters detected using a pan tRNA probe were larger and often more elongated than those detected using the single tRNA probe (**Figure 1D**).

As mRNAs and rRNAs require microtubule (MT)-based transport for their localization in the heart^4^, skeletal muscle^5^, and brain^15–17^, we next asked whether tRNAs colocalized with the microtubule network by combining super-resolution immunofluorescence with tRNA HCR-FISH in primary rat cardiomyocytes (**Figures 2A and S2A**), skeletal myofibers (**Figure 2B**) and neurons (**Figure S2B**). Visual inspection indicated high co-localization of tRNA puncta on microtubules in all cell types. To quantify this co-localization, we assessed the percentage of tRNAs bound to microtubules compared to randomized puncta of the same size and shape. To do this we used a trainable waikato environment for knowledge analysis (Weka) segmentation plugin which allows for unbiased object-based colocalization analysis^18^. We found that tRNAs significantly colocalized with the microtubule network (**Figure 2B**) compared to randomly distributed (R.D.) puncta of the same size and shape. Additionally, tRNAs colocalized with microtubules were larger and had more elongated morphologies compared to tRNAs off the microtubule network in both skeletal and cardiac muscle cells (**Figures 2B and S2A**). Together, these observations indicated that discrete complexes containing multiple tRNAs non-randomly interact with the microtubule network or associated structures.

**Figure 2:**
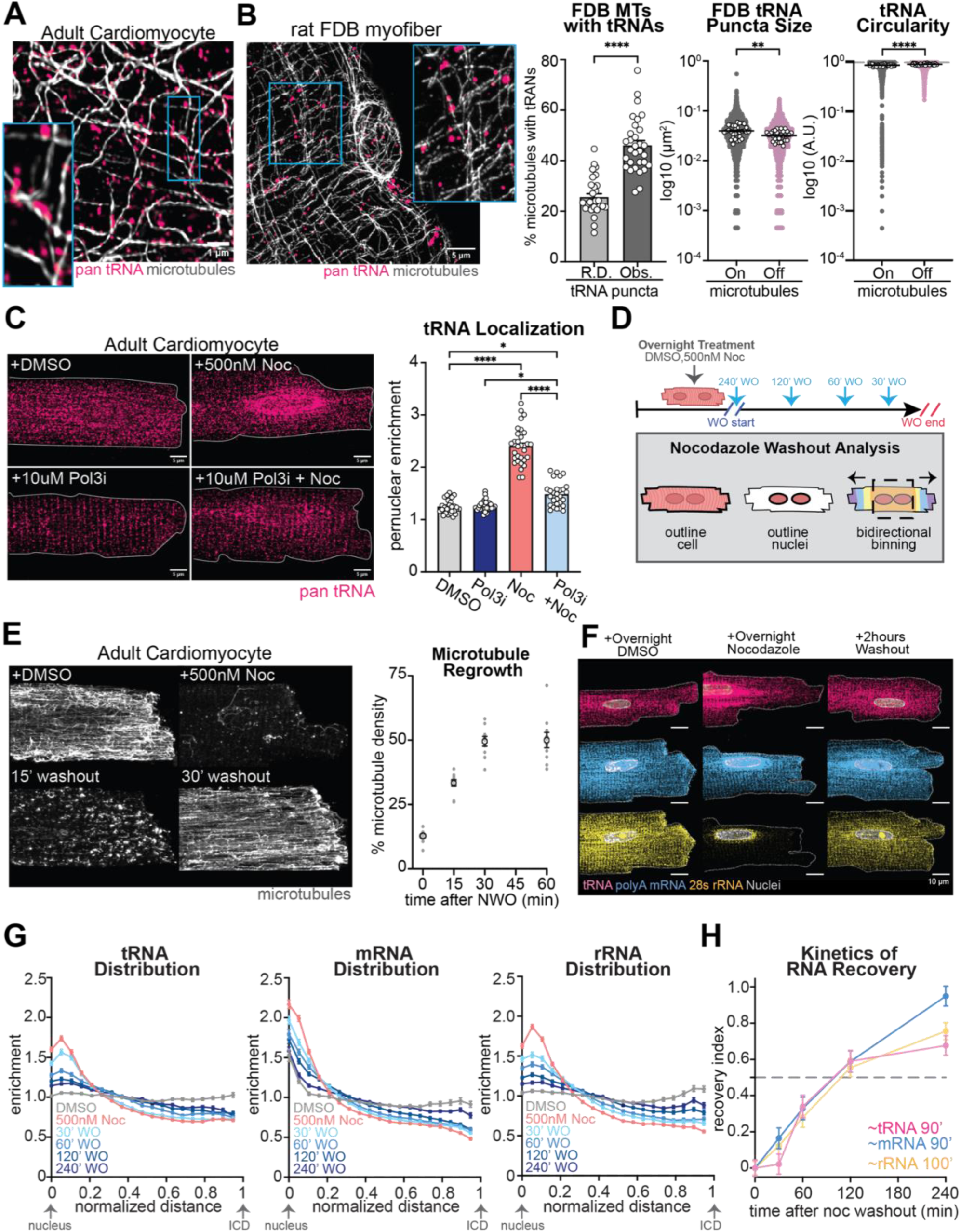
Microtubule-mediated control of tRNA localization. (A) Representative images of microtubules (grey) and pan tRNAs (pink) in ARVMs and FDBs. (B) Quantifications of the observed fraction of microtubules colocalized with pan tRNA (dark grey, Obs.) compared to that expected from random chance using randomly distributed puncta of the same size and shape (light grey, R.D.) and quantification of tRNA puncta size and circularity on (dark grey) or off (pink) microtubules plotted on a log10 scale, with each individual graphical dot indicating an individual puncta measurement and white dots indicating means for each individual cell. (C) Representative images of tRNA localization in ARVMs treated overnight with DMSO (grey), 10uM Polymerase III inhibitor (Pol3i, Blue), 500nM Noc (light red), or a combination of Noc and Pol3i (Pol3i + Noc, light blue) and quantification of enrichment of tRNA signal in the perinuclear region (2 micron ring outside of nucleus) relative to the mean cytosolic signal intensity. (D) Schematic depicting the nocodazole washout assay (described in methods). (E) Representative images of microtubules (grey) in ARVMs treated with DMSO, treated with overnight nocodazole, or at 15 and 30 minutes following nocodazole washout and quantification of microtubule density as a measurement of microtubule regrowth (*n* = 10). (F) Representative images of endogenous tRNAs (pink), polyA mRNA (blue), and 28s rRNA (yellow) following DMSO (left) overnight nocodazole treatment (middle) or 2 hours after washout (right) in ARVMs. (G) Quantifications of tRNA, mRNA, and rRNA enrichment over the cytosolic mean signal intensity as a function of normalized distance from the edge of the nucleus to the intercalated disc (ICD). (H) Quantification of RNA recovery index (the 50% fractional recovery rates) of tRNA (pink), mRNA (blue), and rRNA (yellow) in washout experiments. All experiments were done in biological triplicate (*N* = 3), with 10 cells imaged per biological replicate (*n* = 10), unless noted. Data are presented as the mean ± SEM. *P < 0.05, **P < 0.01, ***P < 0.001, ****P < 0.0001. Two-sided Student’s t-test in experiments from cell averages in b and via one-way ANOVA with post hoc Tukey’s multipe comparisons in experiment c.

To test whether an intact microtubule network is required for the localization of tRNAs, we examined tRNA localization following overnight treatment with a low dose (500 nM) of the microtubule depolymerizing agent nocodazole (Noc). Disruption of the microtubule network resulted in tRNA accumulation in the perinuclear space in cardiomyocytes and myofibers (**Figures 2C, S1D-E, and S2C**). Closer visualization and puncta quantification confirmed displacement of tRNAs from fragmented microtubule tracks, which was concomitant with a reduced size and rounder shape of tRNA puncta (**Figure S2A**). We postulated that perinuclear accumulation of tRNAs upon microtubule disruption could be attributed to 1) a redistribution of pre-existing cytosolic tRNAs, or 2) a failure of newly synthesized tRNAs to be transported away from the nucleus. To discriminate between these two hypotheses, we disrupted microtubules with and without concomitantly blocking new tRNA synthesis with Pol3 inhibition. Short-term Pol3 inhibition prevented the nocodazole-dependent perinuclear accumulation of tRNAs (**Figure 2C**), indicating that the microtubule network is needed for the distribution of newly synthesized tRNAs. Taken together, the above observations support a hypothesis where tRNAs tether to microtubule tracks (likely via a deformable structure given the elongated morphologies observed), which when disrupted limits their distribution throughout the cell and promotes a more spherical, lower energy configuration to tRNA assemblies.

We next took advantage of the reversible nature of nocodazole to examine the kinetics and reversibility of tRNA distribution upon microtubule depolymerization and regrowth (**Figure 2D**). We confirmed that 500 nM nocodazole largely depolymerizes the cardiomyocyte microtubule network and that the network reforms within ∼30 minutes of nocodazole washout (**Figure 2E**). We then fixed cardiomyocytes at different intervals after washout and stained the cells for different RNA species. As expected from past^4^ and current work, each RNA species accumulated around the nucleus following overnight nocodazole treatment and then was redistributed throughout the cell at 2 hours following nocodazole washout (**Figure 2F**).

Since cardiomyocyte microtubules are primarily longitudinally oriented with the minus-ends preferentially localized at the nuclear periphery and plus-ends directed towards the intercalated discs (ICD), we developed an analysis pipeline to quantify tRNA, mRNA, and rRNA enrichment (relative abundance over mean cytosolic abundance) in discrete spatial compartments at normalized distances from the nucleus to the ICD. All 3 RNA species showed reversible microtubule-dependent redistribution throughout the myocyte; however, some subtle differences were observed. Specifically, tRNAs accumulated around the perinuclear space upon microtubule disruption and were distributed throughout the cell following washout, but never recovered enrichment at the ICD. mRNA and rRNA, which exhibit basal enrichment at the Z-disc and ICD, also displayed perinuclear accumulation after microtubule disruption, yet were distributed to the ICD with some delay following MT regrowth (**Figure 2G**). To better characterize the kinetics of RNA dispersal away from the nucleus, a recovery index was calculated from the enrichment curves in **Figure 2G** (see methods for details). The recovery index was normalized (DMSO = 1, nocodazole = 0) and reflects how the RNA species disperse to peripheral regions with time after nocodazole washout, approaching the values observed in DMSO treated cells. While the recovery indices of the RNA species were similar, we noted subtle kinetic differences. The slopes of mRNA and rRNA recovery were more linear in nature, while tRNA recovery was largely absent in the first 30 minutes when the MT network regrew, and then exhibited the most rapid phase (highest slope) of recovery (**Figure 2H**). These data support that pools of tRNAs undergo active microtubule-based transport, as do rRNA and mRNA, but that their transport mechanisms or terminal destinations may be distinct.

### Live-cell single-particle tracking reveals multi-modal tRNA motility

We next aimed to visualize tRNA transport in real-time. Bulk tRNAs can be labeled by attaching a Cy5 fluorophore to the dihydrouridines within their D-loop^19^. Using this method, we delivered bulk Cy5-tRNAs to differentiated C2C12 muscle cells via transient transfection and then performed live-cell imaging to monitor and quantify^20,21^ tRNA movements (**Figures 3A, 3B, and Video S1**). Cy5-labeled tRNAs introduced into myocytes formed discrete, trackable puncta often displaying dynamically changing and elongated morphologies. tRNA puncta exhibited heterogeneous trajectories ranging from static/confined tRNAs (moving no further than 0.1 microns over a 2-minute acquisition), to those undergoing short-distance movements (moving between 0.1 to 1 microns), and a population undergoing long-range (1-10 um) directed transport (**Figures 3B and 3C**). To determine whether tRNA charging was required for long-range transport, we concurrently delivered bacterial and yeast tRNAs to C2C12 myoblast cells, as mammalian cells have their own specific aminoacyl-tRNA synthase enzymes that can charge yeast, but not bacterial, tRNAs^22–25^ (Reviewed in ^26–30^). Both yeast and bacterial labeled tRNAs displayed the ability to undergo long-range transport in myoblasts (**Video S2**), consistent with the idea that aminoacylation is not strictly required for long-range tRNA transport.

**Figure 3:**
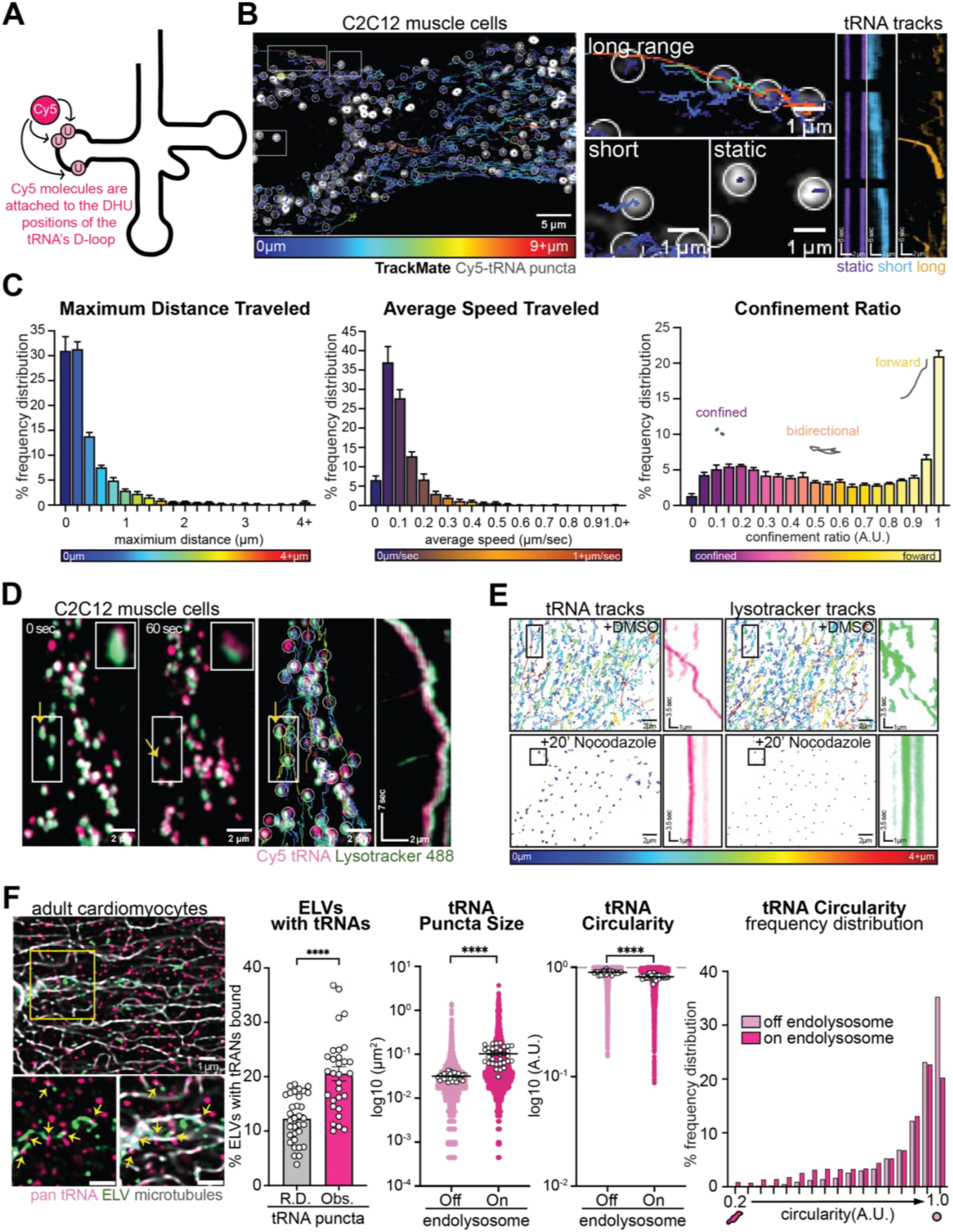
tRNAs undergo long-range active transport by hitchhiking on endolysosomes. (A) Schematic of Cy5-tRNA labeling strategy. (B) Representative snapshot of Cy5-tRNA (white puncta with white circles) in C2C12 cells overlaid with tracks color coded by distance traveled (dark blue, <1 µm; dark red ≥ 9 µm), and zoom in (white boxes) and representative kymographs of tRNAs undergoing long range, short or static movements (purple, static; light blue, short; orange, long). (C) Frequency distributions depicting the maximum distance traveled, average tRNA puncta speed, and confinement ratio per punctum, where 0 is confined to an area, and 1 indicates linear forward movement (*n* = 5 myotubes, with approximately 2000 puncta/video analyzed). Heatmaps below each respective graph denote the color coding of maximum and minimum values. (D) Representative video snapshots of C2C12 myotubes with ELVs labeled with Lyostracker (green) and transfected with Cy5-tRNAs (pink), with a white box indicating the track (yellow arrow) zoomed in for the kymograph. (E) Representative Cy5-tRNA and Lysotracker tracks (colored based on maximum distance traveled per punctum, dark blue, <1 µm; dark red ≥ 9 µm) and kymographs from zoomed in black box in C2C12s treated with DMSO (top panel) or treated with 500nM nocodazole for 1h (bottom panel). Pink kymograph indicates tRNA track, and green indicates an ELV track. (F) Representative images of endogenous LamTOR4-labeled ELVs (green), microtubules (grey), and pan tRNAs (pink) in ARVMS and quantification of the percentage of ELVs colocalized with tRNAs (Obs = observed, bright pink) compared to that expected from random chance using randomly distributed puncta of the same size and shape (grey, R.D.), as well as, quantification of tRNA puncta size and circularity plotted on the log10 scale (with each individual graphical dot indicating an individual puncta measurement and white dots indicating means for each individual cell). From non-transformed circularity measurements, frequency distribution of observed tRNA circularity when tRNA are off (light pink) or on (bright pink) ELVs, with 1 indicating a perfect circle and decreasing values from 1 indicating more elongated morphology. All experiments were done in biological triplicate (*N* = 3), with 10 cells imaged per biological replicate (*n* = 10), unless noted. Data are presented as the mean ± SEM. *P < 0.05, **P < 0.01, ***P < 0.001, ****P < 0.0001. Two-sided Student’s t-test in f.

Quantification of the maximum distance traveled per tRNA punctum revealed that tRNAs occupy multiple motile states, with most punctum being confined or only moving a short distance, while a subset undergo directed long-range transport (**Figure 3C**) consistent with microtubule motor-based trafficking. When examining the average speed traveled per particle, the distribution sharply peaked at low velocities consistent with confined or static particles, with a smaller subset reaching higher velocities (>0.3–0.5 µm/s) within the known range of kinesin/dynein-driven transport^31,32^(**Figure 3C**). This non-normal velocity distribution is inconsistent with Brownian motion, and the coexistence of slow and fast populations suggests state switching between motor binding and unbinding. When examining the frequency distribution of confinement ratios (**Figure 3C**; 0–0.3 is strongly confined, 0.4–0.6 represents bidirectionality and oscillatory behavior, and>0.7 indicates strong forward movements), we observe a dominant immobile/locally confined population, a substantial intermediate population showing short-range bidirectional behavior, and a definitive processive, high-speed, long-distance population indicative of motor-driven transport.

Performing two-color live cell imaging with Cy5-tRNAs and the live cell microtubule dye SpyTubulin we confirmed that tRNAs undergoing long range transport were moving on microtubule tracks in C2C12 myocytes (**Video S3**) and neurons (**Figures S2D and Video S4**). In myofibers exhibiting frequent long range tRNA trajectories, nocodazole treatment subsequently halted all transport processes (**Video S5**). Together, these analyses and real-time observations indicate that tRNAs toggle between mobility states, a hallmark of actively regulated RNA cargo that traffic on microtubules rather than freely diffuse throughout the cell.

### Active transport of tRNAs occurs through endolysosomal hitchhiking

Active transport of mRNA on microtubules is a conserved process, predominantly occurring through the formation of mRNA-RNA binding protein (RNP) interactions that come together to form liquid-liquid phase-separated granules (reviewed in ^17^). In the canonical model of mRNA transport, these RNAs are quiescent until reaching their terminal endpoint, where they are de-repressed and made available for translation. For a smaller percentage of transport-competent RNP granules, they are also capable of tethering (“hitchhiking”) to endolysosomal vesicles (ELVs) to achieve long-range transport, a process that is conserved from fungi to neurons^33–38^.

Based on our observations that microtubule-associated tRNA puncta exhibited elongated morphologies like membrane-bound organelles (**Figure 2A**), we hypothesized that tRNAs may utilize ELV hitchhiking to achieve active transport. To determine if tRNAs co-transported with ELVs, we again performed two-color live cell tracking, this time following the trajectories of Cy5-labeled tRNAs with LysoTracker-labeled ELVs (**Figure 3D**). We observed that tRNAs were specifically co-trafficking on the outside of ELVs in C2C12 muscle cells (**Figure 3D**). tRNA and ELV long distance movements were closely linked, and these directed transport events were almost completely abrogated for both tRNAs and ELVs following 20 minutes of nocodazole treatment (**Figures 3E and Video S6**). We also observed that tRNA-ELV hitchhiking occurs in primary neurons (**Video S7**). To ensure what we were observing was not an artifact derived from potential confounding factors introduced by lipofection (which utilizes an endosomal pathway for nucleotide internalization and escape), we introduced labeled tRNAs into myocytes through a non-endosomal route (electroporation) and found that ELV hitchhiking events occurred in the same fashion independent of the route of exogenous tRNA administration (**Video S8**). Together, our studies suggest that tRNAs may hitchhike on ELVs to achieve microtubule-based transport, rather than independently engaging with MT motors or other RNPs.

We next combined tRNA FISH with immunofluorescence labeling of ELVs in primary adult cardiomyocytes (**Figure 3F**), C2C12 myoblasts, muscle tissue sections, primary adult myofibers, and primary neurons (**Figures S2E-S2G**) to observe endogenous tRNA-ELV interactions. The percentage of tRNAs that colocalized to the outside of ELVs was significantly higher than expected by random chance, as determined by randomly distributing the tRNA puncta across the same ELV images (**Figures 3F and S2F**). Moreover, tRNA puncta on the exterior of ELVs were markedly larger and more elongated than those not bound to ELVs (**Figures 3F and S2F**), supportive of multiple tRNAs associating with ELV membranes in a transport complex. Taken together, results from live and fixed cell experiments support ELVs as active carriers of tRNA cargo, suggesting a new mechanism of tRNA transport.

### Determining the components required for tRNA-ELV hitchhiking

Kinesin-1, encoded by *Kif5b*, is the predominant muscle motor responsible for ELV transport^39–42^ (reviewed in^43^). To test the necessity of Kinesin-1 for tRNA localization, we transduced adult primary cardiomyocytes with a previously validated adenovirus encoding shRNA to knockdown *Kif5b* (shKif5b)^4^. Following 96h of transduction and ∼65% knockdown, we observed a perinuclear accumulation of tRNAs upon *Kif5b* KD (**Figures 4A, S3A, and S3B**). We further examined the role of Kinesin-1 using live-cell tRNA tracking experiments. In C2C12 cells receiving scramble siRNA sequences, we observed tRNAs undergoing long-range transport, yet cells receiving siRNAs targeting Kif5b and achieving ∼50% knockout, we observed tRNAs undergoing short and sparse tRNA movements (**Figures 4B, S3C, and Video S9**). Long-distance transport events (≥ 1.5 microns) were significantly reduced in *Kif5b* depleted myocytes (**Figure 4B**), suggesting that Kinesin-1 mediates tRNA localization and long-range transport in muscle cells.

**Figure 4:**
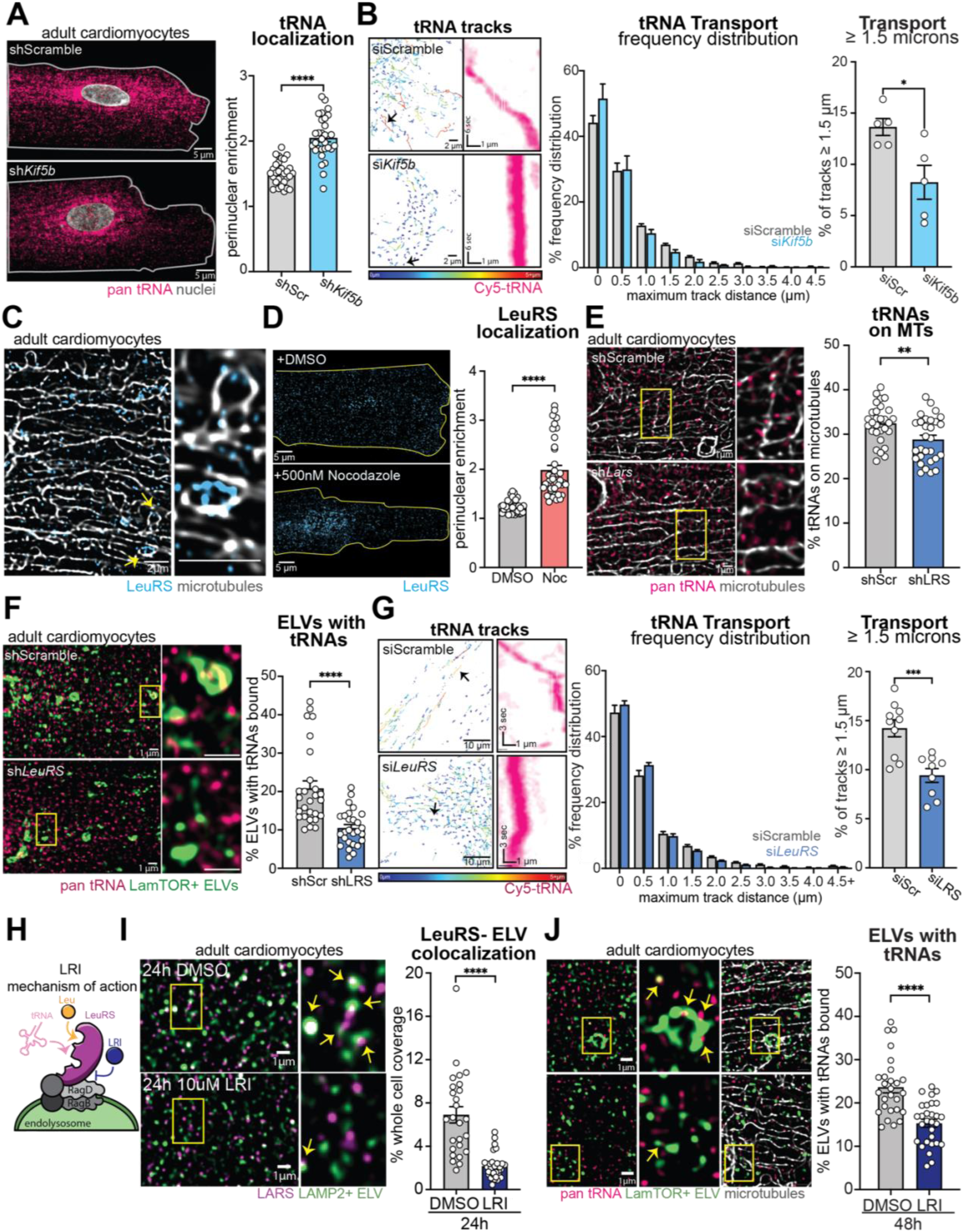
Kinesin-1 and LeuRS mediate tRNA-transport. (A) Representative images of endogenous pan tRNAs (pink) and quantification of perinuclear enrichment relative to cytoplasmic levels in ARVMs infected with an adenovirus encoding short hairpins to knockdown *Kif5b* (Ad-shKif5b, light blue bar) or a scramble (Ad-ShCtrl, light grey bar) control sequence for 96h. (B) Representative Cy5-tRNA tracks (color coded by maximum distance traveled, dark blue, <1 µm; dark red ≥ 5 µm) and representative tRNA kymograph (arrow points to specific track) in myotubes after 72h treatment with 50 nM siRNAs targeting scramble sequences (siScr, grey bar) or siRNAs targeting KIF5b (si*Kif5b*, light blue). Frequency distribution of maximum distance travelled per tRNA punctum and measurements of the percentage of tracks ≥1.5 microns ( *n* = 5 myotubes, with approximately 500 puncta/video analyzed). (C) Representative image of endogenous Leucyl-tRNA synthetase (LeuRS, blue) and microtubules (grey) in an ARVM, with yellow arrows indicating the area of zoom where LeuRS is decorating microtubules. (D) Representative images of LeuRS (blue) localization in ARVMs treated overnight with DMSO (grey bars) or 500nM nocodazole (Noc, red bars) and quantification of perinuclear enrichment relative to cytoplasmic levels. (E) Representative images of pan tRNA (pink) and microtubule in ARVMs treated 96h with an adenovirus encoding short hairpins targeting a scramble sequence (Ad-sh-Scr, grey) or LeuRS (Ad-shLRS, blue) and quantification of the percentage of tRNA puncta colocalizing with microtubule tracks. (F) Representative images of ARVMs transduced with Ad-shScr (shScr) or Ad-shLeuRS (shLRS) after 96h after infection and staining of LamTOR4+ ELVs (green) and pan-tRNA (pink) and quantification of the percent of ELVs colocalized with tRNA puncta. (G) Representative images of Cy5-tRNA tracks (color coded by maximum distance traveled, dark blue, <1 µm; dark red ≥ 5 µm) and representative kymographs (tRNA track, pink) from the selected tRNA puncta (arrow) in C2C12s treated for 96h with 50nM siRNAs to knockdown LeuRS (siLRS, blue) or a scramble sequence (siScr, grey) and frequency distribution of maximum distance travelled by puncta, and measurements of the percentage of tracks ≥1.5 microns ( *n* = 9 myotubes, with approximately 500 puncta/video analyzed). (H) Schematic of how the BC-LI-0186 (LeuRS-RagD inhibitor, LRI), which inhibits LeuRS from binding the RagD site on the outside of endolysosomes (ELVs), functions. (I) Representative images of LeuRS (purple) and ELV (green) staining in ARVMs treated with DMSO or 10 µM LRI for 24 hours (yellow arrows indicate sites of LeuRS-ELV colocalization) and quantification of the abundance of LeuRS-ELV colocalization puncta. (J) Representative images of ARVMs treated for 48h with DMSO or 10 µM LRI and stained for endogenous pan tRNAs (pink), microtubules (silver) and ELVs (green) with yellow box indicating area of zoom, and yellow arrows indicate ELVs with tRNAs on the outside. Quantifications of the percent of ELVs colocalized with tRNA. All experiments were done in biological triplicate (*N* = 3), with 10 cells imaged per biological replicate (*n* = 10), unless noted. Data are presented as the mean ± SEM. *P < 0.05, **P < 0.01, ***P < 0.001, ****P < 0.0001 as determined by two-sided Student’s t-test.

We next sought to identify molecules that may link tRNAs or tRNA binding proteins to ELVs on microtubule tracks. We first screened canonical tRNA binding proteins to ask if their localization was sensitive to microtubule disruption. Neither eEF1a1, which binds and shuttles tRNAs to the ribosome, nor GlyRS, which charges tRNA^Gly^, co-localized with microtubules nor was sensitive to microtubule disruption, despite the fact that the localization of tRNA^Gly^ itself was microtubule-sensitive (**Figures S4A-S4F**). Intriguingly, out of the family of tRNA synthetases, Leucyl-tRNA synthetase (LeuRS) has been shown to uniquely and directly interact with Rag GTPase (RagD) on the outside of ELVs ^44,45^. In support of LeuRS serving as a potential tRNA-ELV adaptor, we found that LeuRS puncta often decorated the microtubule network (**Figure 4C**) and exhibited robust perinuclear accumulation upon microtubule disruption (**Figure 4D**), phenocopying the effect on tRNA localization. To more directly test whether LeuRS mediates tRNA-ELV hitchhiking, we generated an adenovirus encoding shRNA to knockdown LeuRS (shLRS). We confirmed significant but partial (∼50%) knockdown of LeuRS following 96h of AdV-shLRS treatment (**Figure S5A**). Partial depletion of LeuRS reduced the fraction of tRNAs co-localized with microtubules (**Figure 4E**), and the fraction of ELVs co-localized with tRNAs (**Figure 4F**). In live-cell imaging studies, partial LeuRS depletion significantly reduced long-distance tRNA transport (**Figures 4G, S5B, and Video S10**).

To determine whether LeuRS was also required for long-distance tRNA transport in neurons, we first validated robust siRNA depletion of LeuRS in primary cortical neurons (**Figure S5C**). We also validated siRNA depleting GlyRS (SiGRS, **Figure S5D**) to test the specific requirement of LeuRS for tRNA transport. We observed that depletion of LeuRS led to fewer long-distance tRNA transport events and more confined, less bi-directional tRNA movements in neuronal projections, while depletion of GlyRS had no effect on tRNA transport (**Figures S5E-S5G and Video S11)**.

While supportive of a general role in tRNA transport, genetic approaches to deplete LeuRS cannot distinguish between direct disruption of LeuRS-ELV interactions or indirect consequences of LeuRS depletion. To more directly disrupt LeuRS-ELV association without disrupting canonical functions of the tRNA^Leu^ charging enzyme, we took advantage of the small molecule inhibitor BC-LI00186 (abbreviated LRI for <u>L</u>euRS-<u>R</u>agD Inhibitor). LRI prevents LeuRS from binding RagD on ELV membranes but does not impact LeuRS expression or aminoacylation abilities (**Figure 4H**)^46,47^. We first validated its utility in cardiomyocytes and found that LRI treatment effectively displaced LeuRS from ELVs (**Figure 4I**). Consistent with the proposed model of LeuRS as a tRNA-ELV adaptor protein, LRI treatment reduced the number of ELVs associating with tRNAs (**Figure 4J**). Taken together, these data support LeuRS as an adaptor molecule linking tRNAs to ELVs through RagD binding, enabling microtubule motors to transport them as cargo.

### tRNA-ELV hitchhiking supports distributed protein synthesis and maintenance of cell size

As recent reports indicate that mRNA can also hitchhike on ELVs (albeit in RNA granules and via distinct linkers)^6,34,37^, we next asked whether the LeuRS-ELV mechanism was specific for tRNA transport or conserved for mRNA and rRNA distribution. As seen in **Figure 2**, overnight nocodazole treatment leads to perinuclear accumulation of tRNAs, mRNAs, and rRNAs, and the peripheral distribution of each RNA species is largely restored after 4 hours of nocodazole washout and microtubule regrowth. We thus postulated that the addition of 10 μM LRI to the media during nocodazole washout (**Figure 5A**) would 1) block active transport and distribution of newly made tRNAs and 2) could also block active transport and distribution of mRNA and rRNA if they were utilizing this same machinery. As expected, all 3 RNA species accumulated in the perinuclear space upon microtubule disruption and dispersed following Nocodazole washout (**Figure 5B**). However, in the presence of LRI, only the tRNAs remained retained around the nucleus while mRNA and rRNA were clearly distributed to the periphery upon microtubule regrowth. To quantify this RNA retention, we calculated the mean fluorescence intensity of each RNA’s signal in the nuclear and perinuclear compartment relative to the mean cytosolic fluorescence over the time course of the experiment. During the first 30 minutes of washout, when the microtubule network reforms, retention dropped slightly and similarly (some RNA escaped the perinuclear compartment) between all RNA species, independent of treatment condition. However, once the microtubule network was reformed, we observed that all RNA species were effectively cleared from the perinuclear compartment and distributed throughout the cell, except for tRNAs in the presence of LRI, which remained accumulated around the nucleus over the full experimental duration (**Figure 5C**). Further distance mapping of RNA distributions and analysis of recovery kinetics also indicated that LRI uniquely disrupted the ability of tRNA, but not mRNA or rRNA, to distribute throughout the myocyte (**Figures 5D and S6A-6E**). This data indicates that the non-diffusible pool of tRNAs are unable to be redistributed upon LeuRS-RagD disruption, and that LeuRS-ELV mediated RNA hitchhiking is an exclusive mechanism for tRNA distribution.

**Figure 5:**
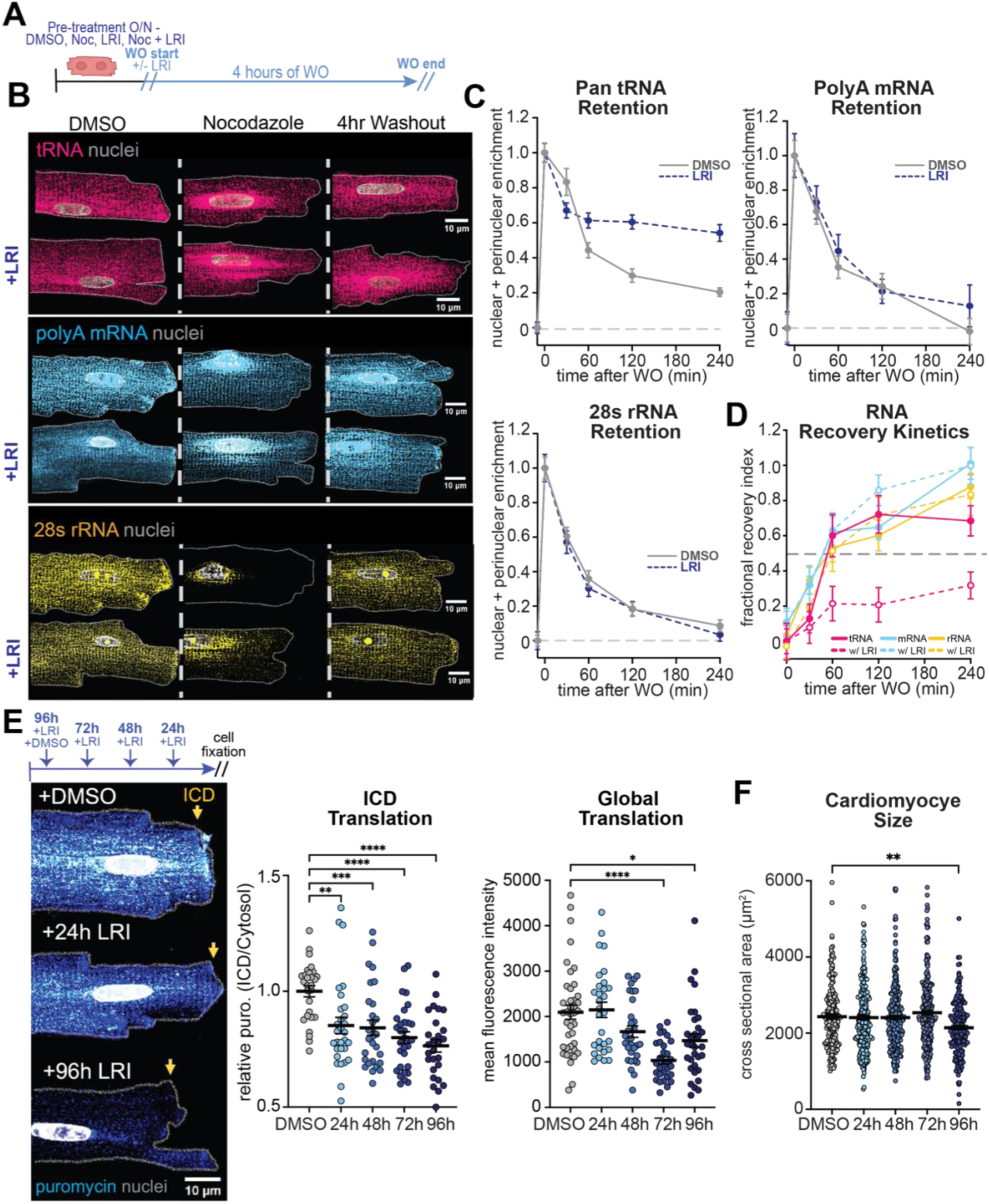
tRNA transport is distinct from messenger and ribosomal RNAs and regulates cardiomyocyte translation and size. (A) Schematic of Nocodazole (noc) washout experiment in the presence of the LeuRS-RagD inhibitor (LRI) or DMSO control. (B) Representative images of pan tRNA (pink), polyA mRNA (blue), and 28s rRNA treated with DMSO overnight (left panel), 500nM nocodazole overnight (middle panel), or following 4 hours of WO (right panel). Top row indicates washout with DMSO, and bottom row indicates washout with 10 µM LRI. (C) Quantifications of combined nuclear and perinuclear enrichment of pan tRNA, polyA mRNA, and 28s rRNA over the course of 4 hours with (dark blue dashed line, LRI) and without (grey line, DMSO) the LRI in washout media. (D) Quantification of RNA recovery index (the 50% fractional recovery rates) of tRNA (pink), mRNA (blue), and rRNA (yellow) with DMSO (solid lines) or LRI (dashed lines) in the media. (E) Experimental schematic and representative images of puromycin (blue, the yellow arrow indicates intercalated disc, ICD) incorporation in ARVMs all fixed at 96h, following treatment with LRI for varying times (24h, 48h, 72h, 96h) and quantification of puromycin-labeled peptides at the ICD relative to cytoplasmic levels and puromycin-labeled peptides throughout the whole cell relative to DMSO control. (F) Measurements of cardiomyocyte area at 96h following exposure to LRI treatment at various timepoints (24,48, 72,96h). All experiments are performed in biological triplicate (*N* = 3), with washout and local translation experiments involving quantification of 10 cells per biological replicate (cell size measurements included more than 50 cells per biological replicate). Data are presented as the mean ± SEM. *P < 0.05, **P < 0.01, ***P < 0.001, ****P < 0.0001, and determined via one-way ANOVA with post hoc Tukey’s multiple comparisons in the experiment.

Having established that tRNA transport can be disrupted without affecting mRNA or rRNA distribution, we next interrogated the impact on cellular protein synthesis. Protein synthesis is distributed throughout the cardiomyocyte, with translation enriched at the level of the sarcomere and intercalated disk^4,48^. Using a puromycin incorporation assay to labeled newly synthesized polypeptide chains^49^, we observed that specifically blocking tRNA-ELV transport lead to a loss of local translation at the intercalated disc within 24 hours (**Figure 5E**). By 72 and 96 hours of treatment, global protein synthesis throughout the cell was also significantly reduced (**Figure 5E**). Consistent with impaired proteostasis, we observed that by 96 hours, LRI-treated cardiomyocytes exhibited reduced cell size compared to DMSO-treated controls (**Figure 5F**). Together, these data indicate that when tRNA transport is specifically disrupted, local and global protein synthesis decline in a time-dependent manner, leading to eventual myocyte atrophy. This data indicates that effective microtubule-dependent tRNA transport is required to support distributed protein synthesis, maintenance of the muscle proteome and cell size.

## Discussion

### tRNAs undergo active transport in mammalian cells

Here, we identify that there are pools of tRNAs that undergo microtubule motor-dependent transport to achieve spatial distribution in mammalian cells. We demonstrate that this process relies on the tRNA-binding protein LeuRS, which mediates the co-transport of tRNAs with endolysosomal vesicles. Consequently, disruption of the microtubule network, Kinesin-1, or this adaptor results in impaired tRNA trafficking, perturbed localized translation, and reductions in cell size. Through selective disruption of LeuRS-ELV interactions, we uncouple the role of tRNA transport from microtubule disruption and show that the mechanism of tRNA-based transport is distinct from that of messenger and ribosomal RNA. We thus establish that tRNAs are actively trafficked cargo that utilize a distinct transport mechanism to spatially regulate protein synthesis.

Our data does not imply that all tRNAs are actively transported, but rather that certain pools can be. In live cell studies we observed (1) a number of static puncta with anchored assemblies, which we suspect correspond to sites of either translation, RNP accumulation, or structural association, (2) a number of short range tracks with jittery, bi-directional transport with particles seemingly corralled in specific locations, and (3) a pool undergoing long-range, directed paths indicative of processive motor-driven movement. The juxtaposition of these differential movement modalities supports the notion that tRNAs likely toggle between motility states, a hallmark of actively regulated RNA cargo. In concurrence with this notion, in our fixed cell studies we identified many smaller, rounder, tRNA puncta that tend not to co-localize with the microtubule network or ELVs. We speculate that these puncta may represent freely diffusing or single tRNA molecules, and it is also possible that some smaller puncta remain unidentified due to resolution limits or signals close to background noise. Consistent with this idea, using STORM microscopy that can detect single fluorophore blinking events at ∼20 nm resolution, we observed a larger population of small tRNA molecules. Nonetheless, across all modalities (STORM and AiryScan imaging of endogenous tRNAs, live-cell imaging of exogenous tRNAs), we observe a pool of larger, elongated tRNA puncta with sizes compatible with multiple tRNAs in complex. These puncta associate preferentially on microtubule tracks and with ELVs, likely as part of a larger transport complex.

### How do tRNAs get directed to ELVs?

The finding that tRNAs associate with ELVs through LeuRS raises several questions pertaining to how tRNAs are directed to these vesicles and the specificity of this adaptor. For mRNA transport, specific association with RBPs often drives the separation of these transcripts into granules, which are then targeted for transport by cytoskeletal motor proteins (reviewed in ^50^). mRNA localization into transport competent granules is believed to occur through: 1) RBP recognition of short “zip code” sequences within the 3’ and 5’ untranslated regions (UTRs) and subsequent directing of those mRNAs to specific locations (reviewed in ^7,51,52^); 2) RBP recognition of complex secondary structures ^9,53,54^ and chemical modifications to specific nucleosides^55–58^. Considering that tRNAs lack 3’ and 5’ UTRs, it seems less likely that RBPs identify tRNAs through a zip code motif, but perhaps through their unique secondary structure. This concept gains some support by the identification of LeuRS as an adaptor linking tRNAs to ELVs. LeuRS canonically functions as a tRNA synthetase, meaning it has ATP-dependent catalytic activity that charges amino acids to its cognate tRNA and an editing activity that ensures the correct amino acid is loaded and charged. However, to bind tRNA, it recognizes the long variable arm and discriminator base A73 of tRNA^Leu^ prior to charging^59^. Indeed, all class 1 aminoacyl tRNA synthetases (aaRS), of which LeuRS belongs, primarily bind tRNAs through interactions with the minor groove of the tRNA acceptor stem and hence recognize tRNA structure^60^. This means they can also bind non-cognate tRNAs, but at a lower affinity, with that low affinity increasing depending on structural similarities to tRNA^Leu 61,62^. Considering that preventing LeuRS-ELV interactions was sufficient to disrupt the majority of tRNA hitchhiking, this could be reflective of LeuRS’ ability to bind non-cognate tRNAs based on structural engagement^63^, but not charge them^64^, a model supportive of a noncanonical role for LeuRS as a tRNA transport adaptor. And, considering that LeuRS is also part of the multi-tRNA synthetase complex ^65^, this raises the question of whether LeuRS, of all the 20 aaRS enzymes, is the sole mediator of tRNA-ELV hitchhiking, or if others could also participate. This assigns a previously unrecognized, non-canonical function to an aminoacyl-tRNA synthetase, expanding the roles of ARS proteins beyond aminoacylation and signaling^66^ to also include RNA spatial organization.

### The conservation of tRNA transport

Questions remain about whether this process is conserved in different cell types of varying complexity, particularly when considering that tRNAs and the aminoacyl-tRNA synthetase enzymes (aaRS) which charge them have diverged in sequence throughout evolution (Reviewed in ^67^). Neurons, skeletal muscle myofibers, and cardiomyocytes represent highly polarized, large mammalian cells that are largely incapable of division. These cells demonstrate how active transport of RNAs is needed to support efficient local translation over long distances or diffusion restricted environments^4,5^. Intriguingly, while not demonstrating active transport, prior attempts to assess tRNA dynamics in live mouse embryonic fibroblast cells^9^ suggested these molecules exhibited distinct, non-linear cytoplasmic distributions inconsistent with molecular diffusion. Thus, it remains unclear whether active tRNA transport is conserved beyond the complex cell types examined here. To this end, we initially tested the transport competence of yeast and bacterial tRNAs, assuming that if this were an evolutionary adaptation, more primitive bacterial tRNAs may not be transport-competent. This experiment also informs about the charging status of these moving tRNA pools, as bacterial tRNAs cannot be efficiently charged in mammalian cells due to species-specific tRNA synthetases^22–25^ (Reviewed in^26–30^). Interestingly, both yeast and bacterial tRNA types, despite considerable sequence variation and charging capabilities, but conserved secondary structure, were capable of transport in C2C12 myoblasts and myotubes. We thus speculate that tRNA structure, and not sequence motifs nor charging status, may be important for recognition by mammalian ELV transport complexes, representing a potential evolutionarily repurposed mechanism for supplying distal translation zones in large or structurally dynamic cells.

### Transport-competent tRNAs as drivers of cell growth

Our findings highlight and motivate further inquiry into the physiological consequences of disrupting tRNA transport, as well as the broader role on ELVs in spatial proteome regulation. These organelles, once viewed as the “garbage disposals” of cells, are now recognized as signaling hubs and long-range trafficking platforms capable of supporting localized translation. For instance, some ELVs have been shown to transport mRNAs that regulate ELV fusion, suggesting that spatiotemporal coupling of mRNA transport and translation may facilitate the local production of proteins that modulate the functionality of ELVs themselves^68^. Moreover, mRNA-ELV hitching has been demonstrated to be important for the transport of transcripts that encode key genes involved in maintaining mitochondrial homeostasis and preventing axonal degeneration ^6,34^. In muscle cells, here we find that disrupting ELV-mediated tRNA localization seems to affect translation initially at the distal ends of these long, polarized cells. Subsequently, over the time course of tRNA turnover, we observed a gradual reduction in global protein synthesis, and ultimately a loss of myocyte size. Given that hypertrophic stimuli elevates polymerase III activity, which expands the tRNA pool size (reviewed in ^69^), these findings raise the possibility that tRNA-mediated regulation of translation may be an underappreciated determinant of cellular proteostasis and growth.

### Concluding Remarks

We present here initial evidence that tRNAs undergo directed long-range transport along microtubule tracks. This is primarily achieved through endolysosomal hitchhiking, a process adjacent to yet complementary with RNP trafficking. We propose that LeuRS tethers tRNAs to endolysosomes, which engage with microtubule motors to drive their long-range transport. Inhibition of the tRNA-LeuRS-ELV interface disrupts tRNA, but not mRNA or rRNA transport, and is sufficient to collapse local translation, indicating the importance of correctly localizing the tRNA pool for the maintenance of protein synthesis and cellular size.

## Materials and Methods

### Ethics declarations

All presented experiments comply with the standards set forth by the Institutional Animal Care and Use Committee at the University of Pennsylvania and the Guide and Care and Use of Laboratory Animals published by the US National Institute of Health. All protocols are approved by the University of Pennsylvania Institutional Animal Care and Use Committee and Institutional Biosafety Committee.

### Animals

Male and female C57BL6/J mice and Sprague Dawley rats up to 24 weeks of age were used in this study. Animals were housed at 72° Fahrenheit under a 12-h light/12-h dark cycle and maintained on a standard chow diet. Animals had ad libitum access to food and water.

### Primary adult rat ventricular myocyte (ARVM) isolation

ARVMs were isolated from 8 to 16-week-old Sprague Dawley rats using Langendorff retrograde aortic perfusion with an enzymatic solution as previously describe^4^. Briefly, the heart was removed from an anesthetized rodent under isoflurane and retrograde-perfused on a Langendorff apparatus with a collagenase solution. The digested heart was then minced and triturated with glass pipettes to free individual cardiomyocytes. The resulting supernatant was separated and centrifuged at 300 rpm to isolate cardiomyocytes. These cardiomyocytes were then resuspended in ARVM media (Medium 199 (Thermo Fisher) supplemented with 1x insulin-transferrin-selenium-X (ITS, Gibco), 1 μg/μL primocin (InvivoGen), 20 mM HEPES, pH = 7.4 (UPenn Cell Center), and 25 μM of cytochalasin D (Cayman Chemical)) at low density and cultured at 37 °° C and 5% CO_2_.

### Primary rat FDB isolation

Flexor digitorum brevis (FDB) muscles were harvested bilaterally from 8 to 12-week old Sprague Dawley rats following euthanasia. To obtain single myofibers by enzymatic dissociation, FDB muscles were isolated, dissociated for 3-5 hours at 37 °C, and cultured in a humidified incubator at 37 °C (5% CO2) as previously described^70–72^. FDB muscles were dissected and digested in Dulbecco’s modified Eagle’s medium (Thermo Fisher) and 4 mg/ml collagenase (Type I #C0130, Sigma) for 3 h at 37 °C. Muscle fibers were then plated on glass-bottomed culture dishes (# P35G-1.0-14-C, Matek Inc) coated with laminin (#23017015, Life Technologies). After plating, cultures were incubated in DMEM, 10% fetal bovine serum (FBS, #10100139, Life Technologies) and 0.1% gentamicin (#15710064, Life Technologies,) for treatment overnight with DMSO or 500 nM Nocodazole.

### Primary postnatal cortical neuron isolation

Primary cortical neurons were isolated from P0-P3 pups from C57/BL6 mice. Cortical tissue was isolated in cold 1x Hibernate-A (Gibco #A12475-01) and cut in to small pieces (∼2mm) under sterile conditions. Cortices were pooled together and dissociated into single cells via incubation in 20U/mL Papain plus DNase (Worthington Biochemical Corporation, #lk003178 and #LK003172) for 15-20 minutes at 37°C. Subsequently, they were manually dissociated by trituration with a 5 ml Serological pipette 3-4 times followed by 25-30 times with a p1000 pipette tip. Cells were centrifuged at 300 x g for 5 min and washed with 1x PBS (with calcium/magnesium). Pellets were then resuspended in Neurobasal-A (Gibco) supplemented with 1% B27 Plus (Gibco), 1% MEM-NEAA (Gibco), 1% GlutaMax (Gibco), 1% Penicillin/Streptomycin (Gibco), 1% Heat-Inactivated Fetal Bovine Serum (Gibco), and 5ug/mL Laminin (Sigma). Live cells were counted using trypan blue and plated (300,000 cells/mL) onto Mattek Dishes. Plates were coated with 50ug/mL poly-D-lysine (Sigma #P1149) in 0.1M Borate Buffer pH 8.5, washed 3 times in sterile H2O and dried for at least 1hr in the TC hood or coated with Matrigel (Corning). Cells were incubated humidified environment at 37°C in 5% CO2, and the following day media was changed to maintenance media without 1% Heat-inactivated FBS. 50% Media was subsequently changed twice a week until the experimental endpoint.

### C2C12 myocyte cell culture

Myoblast C2C12 cells (ATCC) were cultured in standard culture medium (20% fetal bovine serum, high-glucose DMEM, 1% Pen-Strep) and differentiated in differentiation medium with daily media changes to promote formation of myotubes at 70% confluence (1%horse serum (Gibco), DMEM high-glucose (Gibco), 1% Pen-Strep (Invitrogen), Insulin-Transferrin-Selenium Supplement (ITS, Gibco, 100x stock)).

### Preparation of labeled tRNAs

To generate Cy3 or Cy5-labeled tRNAs, bulk *Escherichia coli* and yeast tRNA were purchased from Roche, reduced at dihydrouridine residues, and fluorescently labeled with hydrazide-conjugated dyes (Cy3, Cy5) as previously described19,73-76.

### Transient transfection, Delivery of Cy5-tRNAs, and live-cell labels

For delivery of Cy3 or Cy5-labeled tRNAs (500ng/mL) or siRNAs (50nM, Silencer Select, ThermoFisher, # AM16708, targeting 3 regions of *LeuRS*, *Kif5b*, *GlyRS*, and scramble sequences-pooled siRNAs per target), cells were transiently transfected using Lipofectamine RNAi MAX (Invitrogen, #13778150) according to manufacturer’s instructions. For delivery of labeled tRNAs without the use of lipofectamine, C2C12 cells were electroporated using the Gene Pulser II (Bio-Rad) and Cell Line Nucleofector Kit (Lonza) according to manufacturer’s instructions. To label microtubules of live cells, Spy555-Tubulin (SpiroChrome) was added to the media overnight at a concentration of 1:1000. For labeling of ELVs, 50nM LysoTracker-Green-26 (Invitrogen) was added to the media of cells, an hour being imaging. All imaging experiments were done in phenol-free media.

### Puromycin incorporation in isolated adult rat ventricular cardiomyocytes

To assess the synthesis of new proteins, ARVMS were treated with puromycin (2 µM) for 10 minutes and at 37°C in incubator. *<u>In vitro pharmaceuticals</u>*: Nocodazole (500 nM in DMSO, Thermo Fisher), Puromycin (2uM, A Sigma-Aldrich), Blebbistatin (5 µM in DMSO, Cayman Chemical), Pol3i inhibitor CAS 577784-91-9 (10uM, CAS 577784-91-9, Sigma Aldrich), Lars-RagD inhibitor BC-LI-0186 (10uM, LRI, MedChem Express).

### Virus generation, vectors, and treatment

The adenovirus encoding short hairpins (sh) targeting Kif5b was generated in house 18. The shKif5b construct was directed toward a single target site under the U6 promoter. It was generated and produced by inserting appropriate cDNAs into pENTR for further Gateway recombination in adenoviral expression plasmids. Constructs were then transferred by Gateway recombinase into adenoviral expression plasmid pAdCMV/V5/DEST (Invitrogen). Target site for shKif5b: gcacacagactgagagcaaca. eBFP2 (enhanced blue variant of green fluorescent protein) was used as a marker of transduction for shKif5b. The adenovirus encoding shScramble (no fluorophore) was purchased from Vector Biolabs. The adenovirus encoding short hairpins (sh) targeting LeuRS was generated by Vector Biolabs, as was an shScramble control. The Ad-CFP-U6-r-LARS-shRNA virus (shLeuRS, modified from ADV-292615) construct was directed toward a single target site under the U6 promoter. CFP was used as a marker of transduction. Target site for shLeuRS 5’-CACCGCCGGTTGTGAAGGAGTTAATCTCGAGATTAACTCCTTCACAACCGGC TTTTT-3’. For viruses generated in house, recombinant adenoviral vectors were produced and amplified in HEK 293A cells according to the manufacturer’s protocol (ViraPower Adenoviral Expression System; Invitrogen). Viruses were collected by CsCl gradient centrifugation and dialyzed against a 5% sucrose buffer as the final step. For knockdown of Kif5b in ARVMs, cells were treated with either 0.1 µl/ml shScram or 0.1 µl/ml shKif5b immediately after isolation for 96h. For knockdown of LeuRS in ARVMs, cells were treated with either 1 µl/ml shScram or 1 µl/ml shLeuRS immediately after isolation for 96h.

### Cell fixation for immunofluorescence and HCR-FISH

Isolated ARVMs and primary FDB myofibers were attached to MyoTak-coated(IonOptix, for myocytes) coverslips or dishes. Primary adult mouse neurons were attached to laminin-coated (for neurons) glass dishes. Following time in culture, cells were washed once in PBS, fixed with 4% paraformaldehyde, permeabilized in 0.1% saponin, and stored in 70% EtOH at 4 °C until staining.

### Immunostaining

Fixed samples were brought to room temperature, washed with 1xPBS and blocked (Antibody blocking buffer, Molecular Instruments) for 60 minutes at room temperature. Samples were incubated in primary antibodies overnight at RT (see supplementary table for concentrations, specificity, and catalogue number). The following day, samples were then washed in 4× in 1xPBS with agitation and incubated in a secondary antibodies (1:250) in 50% antibody blocking buffer, 50% 1xPBS overnight. Secondaries used included goat anti-rabbit Alexa Fluor 647, goat anti-mouse Alexa Fluor 647, goat anti-rabbit Alexa Fluor 488, goat anti-mouse Alexa Fluor 488, donkey anti-rabbit Alexa Fluor 555, donkey anti-mouse Alexa Fluor 555 all from ThermoFisher Scientific. For experiments coupling IF with HCR-FISH to detect endogenous tRNAs, probes were added after the washing of secondary antibodies. For those just performing immunostaining, samples were then washed 4×1xPBS with rocker agitation, incubated with Hoechst for nuclear labeling, and mounted on #1.5 coverslips in Prolong Diamond Antifade Mountant (ThermoFisher) for imaging.

### Hairpin chain reaction (HCR) fluorescence in situ hybridization (FISH) detection of tRNA species

Custom hairpin chain reaction (HCR)-FISH (molecular instruments) probe sets were generated for mouse and rat single and pan tRNA probes. For single tRNA species, for each tRNA 2, 18 nucleotide probes were generated targeting positions 1-42 of a given tRNA, with a 2-4 nucleotide gap between them so that they can form an initiator. The same approach was done for the generation of the pan probe, targeting the 50 most abundant cardiac tRNAs as determined from previously published QuantM-tRNA sequencing data 46. During amplification of single tRNA species and pan tRNA ones, HCR hairpin pairs tagged by fluorescent dyes hybridize to each other and initiate the self-assembly that leads to amplification of their signal. For the pan probe, hairpins targeting each of the 50 tRNAs is tagged by the same dye, leading to visualization of multiple species at once. To execute HCR-FISH labeling to tRNAs, the Molecular Instruments mammalian cells protocol was loosely followed. Coverslips were first submerged in Molecular Instruments Hybridization Buffer for 30 min at 37°C in the incubator. Samples were then incubated in probe solution overnight at 37°C in the incubator (all probes added at 1:100 concentration from manufacturer stock solution 200 pmol scale, see below for additional probe set details). The next day, excess probes were removed by washing four times for 5 min in pre-warmed Molecular Instruments Probe Wash Buffer at 37°C with rocking. Samples were washed twice for 5 min in room temperature 5X SSCT and then incubated in room temperature Molecular Instruments Amplification Buffer for 30 min at room temperature. During this time, the appropriate hairpins were snap-cooled using a thermocycler (95°C for 90 seconds, then cooled for 30 min at room temperature, protected from light). Hairpins were added to amplification buffer (all probes added at 1:50 concentration from manufacturer stock solution 600 pmol scale) and added to cells for 2.5 hours at room temperature. After amplifications, cells were washed three times for 5 min in 5X SSCT, then for 10 min where 1:1000 Hoechst was added. Cells were washed once more with 5X SSCT and then mounted on glass slides using Prolong Diamond Antifade Mountant (Thermo Fisher).

### Fluorescence in situ hybridization (FISH) in isolated adult cardiomyocytes to label mRNA and rRNA

For polyA FISH, we used a dT 30-mer 3′ AF 647 (IDT) at a final concentration of 500 nM. For 18S rRNA The Stellaris RNA FISH Protocol for Adherent Cell Types (Biosearch Technologies) was followed as previously described ^4^. Stellaris FISH buffers (Biosearch Technologies) were used for all experiments. Wash Buffer A and Hybridization Buffer were made fresh for each experiment with a new aliquot of deionized formamide. RNAse-free solutions were used whenever possible. Humidity chambers (damp paper towels covered with parafilm) were assembled in 6-well culture plates and coverslips were placed cell-side down on top of 100 mL Hybridization buffer containing the probe set(s) of interest. Coverslips were incubated in the dark at 37 °C for ∼16-20 h. After the incubation, coverslips were transferred to freshly-made Wash Buffer A and incubated in the dark at 37 °C for 30 min. New Wash Buffer A was added again for 30 min, with 1:1000 Hoechst stain added for the last 10 min. Coverslips underwent a final incubation in Wash Buffer B for 5 min at room temperature before mounting on glass slides using Prolong Diamond Antifade Mountant (Thermo Fisher).

### Combination immunofluorescence/HCR-FISH in isolated primary cells

Immunofluorescence was performed prior to FISH, with only a few changes from the standard immunofluorescence-only protocol. Briefly, all reagents were prepared to be RNase-free, either through direct purchase or the usage of DEPC-treated water. Cells were fixed using 4% paraformaldehyde (Electron Microscopy Sciences) in RNAse-free PBS (Gibco) for 10 min, washed two times with RNAse-free PBS, permeabilized with 0.1% Saponin in RNAse free PBS for 10 min and blocked using Molecular Instruments Antibody Buffer for 1 h at room temperature with rocking. Primaries were added in fresh Molecular Instruments Antibody Buffer, and secondaries were diluted in RNAse-free PBS and washed with RNAse-free PBST. After the secondary antibody incubation n 50% antibody blocking buffer, cells were washed, cells in RNAse-free PBS for 10 min with rocking, prior to proceeding to FISH portions of the protocol.

### Airyscan Confocal Imaging

Airyscan confocal imaging for all experiments ARVMs was carried out on either the Zeiss 880 or Zeiss 980 Airyscan confocal microscope operating on an Axiovert Z1 inverted microscope equipped with EC-Plan-Neofluar ×10 air 0.30 numerical aperature (NA), Plan-Apochromat ×20 air 0.8 NA, C-Apochromat ×40 water 1.2 NA, and Plan-Apochromat ×63 oil 1.4 NA objectives. Image analysis was performed using ZEN Black software (Carl Zeiss AG) for Airyscan processing, which involves signal integration from the 32 separate sub-resolution detectors in the Airyscan detector and subsequent deconvolution of this integrated signal. Image processing and analysis was performed using FIJI. <u>For experiments measuring whole cell mean fluorescence intensity</u> using the 63x Objective, z-stacks (6 slices at 1 μm step size) were acquired. For analysis, maximum intensity projections were generated from the full z-stack. <u>For experiments requiring Weka segmentation to look at particle</u> <u>size, shape and colocalization</u>, images were taken with the 63x Objective, at 4x Nyquist sampling and 5× zoom, with 3-5 slice (0.2 microns/slice) z-stacks. Following acquisition, images were Airyscan processed and then deconvolved using Zen Black software (Zeiss, Version 14.0.15.201) using a point-spread-function–based algorithm to reassign out-of-focus signal and improve resolution and signal-to-noise while preserving quantitative intensity. Images were then deconvolution was performed using Zen Blue software (Zeiss, Version 2.5). For all analysis, max-intensity projections were generated from the full z-stack. For <u>images used to calculate cell size</u>, where >50 cells were analyzed per replicate, these were obtained with a 20X 0.8 NA air objective. Z-stacks consisting of 4 optical slices at 2 um intervals (6 um total) were imaged across 25 tiles (5×5), resulting in an image ∼ 1.15 mm x 1.15 mm. The focal plane was adjusted at 16 support points (4×4 grid) throughout the tilescan to correct for potential focal drift. A digital zoom of 1.7× was used resulting in a pixel size of 62 x 62 nm. The images were acquired on the 4Y Fast Airyscan mode. Images were Airyscan processed and then stitched together using the Zen software (Carl Zeiss AG). <u>Live cell experiments</u>were performed with 40× water objective, using an onstage incubator at frame rates with signal averaging denoted in video legends. Videos were acquired on the Zeiss 980. In brief for live cell experiments, cells were imaged for 1-2 minutes per video, at between 2-5 FPS, depending on 1 or 2 color imaging requirements and microscope capabilities.

### Weka quantification of colocalization, puncta size, and shape

Following acquisition of images using the 63× 1.4NA oil immersion objective at 4X Nyquist sampling on the Zeiss 980, images were Airyscan processed, deconvolved, and max intensity projections were made. Masks of each channel of interest were generated by first pulling the single channel into the trainable Weka segmentation plugin in Fiji, training the algorithm to identify structures of interest (tRNA puncta for example) or background, and creating a probability map using the Weka segmentation algorithm. A threshold was applied to the probability maps to generate binary mask images. Masks of interest (tRNA puncta, ELV puncta, MTs) were then used with the analyze particles function in FIJI to identify individual particles. Then, by overlapping particles and masks, percentages of particle colocalization could be determined, where a mean intensity greater than 0 indicates some degree of colocalization. Particle analysis was also used to generate size and shape data. tRNA hitchhiking is defined as the percentage of tRNAs that have any overlap with ELV masks. tRNAs on microtubules is defined as the percent of tRNAs overlapping with microtubule masks. ELV translation sites are defined as the percent of puromycin puncta having any overlap with ELV masks.

### Nocodazole washout experiments and analysis

Cardiomyocytes were treated overnight with the reversible microtubule depolymerizing agent, Nocodazole (500nM). The following day, cells were fixed at different time points after exchanging the media to remove Nocodazole and allow for microtubule regrowth. Images of cardiomyocytes labeled for tRNA, poly A (mRNA), 28S rRNA, and nuclei (Hoechst) were obtained at the different time points following Nocodazole washout. In a separate batch of cells, cardiomyocytes were fixed and labeled for tubulin to quantify the rate of microtubule regrowth. Three channels (RNA species 1, RNA species 2, Hoechst), 7 x 1 μm z-stack images were obtained on the Zeiss 980 or Airyscan microscope using a 63× 1.4NA oil immersion objective in fast mode and Airyscan processed. Maximum intensity projections were used for subsequent analysis. To quantify the density of microtubules, the microtubule images were background subtracted and then thresholded to generate a mask image of microtubules. The cellular microtubule density was measured by quantifying the percent area coverage of the microtubule mask within the cardiomyocyte. To quantify the change in localization of the RNA species, a distance analysis was developed using FIJI (NIH) and MATLAB (v R2023b, Mathworks). In FIJI, the Hoechst channel was used to generate nuclear ROIs that defined the nuclear edges. Straight lines were drawn at the outer nuclear edges that were parallel to the opposing intercalated discs. A distance transform was performed to generate a distance map of the cell with pixel values representing the shortest Euclidean distance from the lateral edge of the nearest nucleus. A cell mask was generated to remove background pixels. Finally, background subtraction was performed on the RNA species channels and exported for downstream analysis. The RNA image, distance map, and cell mask were imported into MATLAB and the fluorescence intensity of the RNA signal from each pixel was binned based on the distance value of that pixel from the nearest nucleus. The distance was normalized to the maximal distance between the edge of the nucleus to the longitudinal edge of the cardiomyocyte (Figure 2 washout analysis) or as concentric rings away from nucleus to ends of cells (Figure 5 washout analysis). Enrichment values were calculated by dividing the sum of the signal intensity from each bin by the average signal intensity of all of the bins. A cumulative distribution function (ie. the integrated RNA fluorescence intensity as a function of distance away from the nucleus to the edge of the cell) was generated. To quantify the dispersion of the RNA signal from the nucleus, the distance where the cumulative distribution function reaches 50% of the total fluorescence level (Dist50) was calculated for each cell. This Dist50 value would be relatively large in the DMSO treated group, relatively small upon nocodazole treatment, and have intermediate values following nocodazol washout. To calculate the recovery index, the Dist50 values were normalized so that the mean values from the DMSO group = 1 and the Nocodazole group = 0.

### TrackMate analysis of particle speed and distances traveled

tRNA particle average speed and maximum distance traveled was determined using the particle tracking software TrackMate^20,21^. In brief, after the acquisition of time-series images, files were imported into ImageJ, and the localization, detection, and subsequent identification of single particles frame by frame was determined by the software for a given video. Single particles were linked to building trajectories over time (single-particle tracking) and the resulting trajectories and links were analyzed after the tracking step to characterize the type of motion. Several parameters (average speed, maximum distance traveled, confinement ratio) were calculated to quantify trajectories.

### Quantification of perinuclear, nuclear and intercalated disc localization

#### Maximum intensity projections were prepared from the 63× images

Preliminary regions of interests (ROIs) were obtained by generating masks of the nucleus, perinucleus (2 μm dilated ring around the nucleus, but not including it), the cytoplasm (the area excluding the nuclear and perinuclear region), and the intercalated disc (2 μm expanded ends of the cell). The mean fluorescence intensities were measured from each ROI. Enrichment was measured by normalizing the ROI mean fluorescence to the mean fluorescence from the cytoplasm. <u>Quantification of 20× images</u>. Maximum intensity projections were prepared from the tilescan images. Preliminary regions of interests (ROIs) were obtained by segmenting tilescan images of ARVM with automated thresholding in FIJI. The identified ROIs were quality controlled by separating fused ROIs from multiple ARVM, excluding ROIs originating from dead cells or debris, and removing ROIs from overlapping ARVM. The mean fluorescence intensities from the remaining ROIs were measured from the maximum intensity projections. The measured cellular fluorescence was normalized to the average fluorescence from the DMSO group or listed as raw values as described in figure legends.

### Adult rat ventricular cardiomyocyte cell size measurement

Cardiomyocyte area was quantified using a custom ImageJ macro that applies Gaussian smoothing, automated thresholding, and size-filtered particle analysis to generate cell-specific ROIs from a high-contrast Hoechst channel optimized for segmentation. Each generated ROIs was then used for area and shape measurements of the total cell.

### STORM Imaging & Analysis

Single-Molecule Localization Microscopy imaging was performed on a Nanoimager system (ONI) equipped with 405-, 488-, 561- and 640-nm lasers, 498–551- and 576–620-nm band-pass filters in channel 1, 666–705-nm band-pass filters in channel 2, a 100X 1.45 NA oil immersion objective (Olympus), and a Hamamatsu Flash 4 V3 sCMOS camera using HiLo illumination. Samples were imaged in fresh βME buffer (1% GLOX (14mg glucose oxidase, 20 mg/mL catalase, 10mM Tris pH 8.0, 50mM NaCl),1% beta mercaptoethanol (βME, 14.3M pure stock solution), and 10% glucose in 50 mM Tris pH8, 10 mM NaCl buffer and deionized water). The Nanoimager system was pre-warmed to 30 C to ensure consistent temperature and reduce drift. Images were acquired at 15 ms exposure for 50,000 frames using 160 mW 640-nm laser illumination. STORM localizations were exported from the Nanoimager software for downstream analysis in MATLAB. Localizations were exported using the following parameters: minimum photon count of 1300, X/Y localization precision of 30 nm, and sigma X/Y below 250 nm. Downstream analysis was conducted in MATLAB (v R2022b, Mathworks) using custom-made code^77^. First, individual cells were cropped. Next localizations from each cell were clustered using DBSCAN analysis^78^ with a search radius of 23.4 nm and minimum number of localizations of 5. For consistent comparison, these parameters were used to analyze every cell in every condition. RNA clusters were then analyzed for area, circularity, and aspect ratio.

### Western Blot

Protein extracts from whole skeletal muscles were generated using RIPA buffer (150 mM NaCl, 1% nonidet P-40, 0.5% sodium deoxycholate, 0.1% SDS, 25 mM Tris pH 7.4), centrifuged (4 °C × 12,000RPM × 20 min), and quantified using the Bio-Rad protein assay (Bio-Rad, # 5000001) as previously described^79^. Standard Western blotting analysis was performed using 4-20% SDS-PAGE gels with the following primary antibodies: GAPDH, LeuRS, and GlyRS. Secondary antibody incubations were done at room temperature for 120 min using LICOR anti-rabbit and anti-mouse secondary antibodies (1:10,000) and then imaged using a LICOR system (LICORbio).

### Statistics and data handling

Statistical analysis was performed using Prism (Version 10.0.2). Statistical test and information on biological and technical replicates can be found in the figure legends. For plots, the mean line is shown, with whiskers denoting standard error of the mean (SEM). Statistical tests for each comparison are denoted in the figure legends. Figures and schematics were generated in adobe illustrator.

### Antibodies and concentrations for IF and IF-HCR-FISH experiments

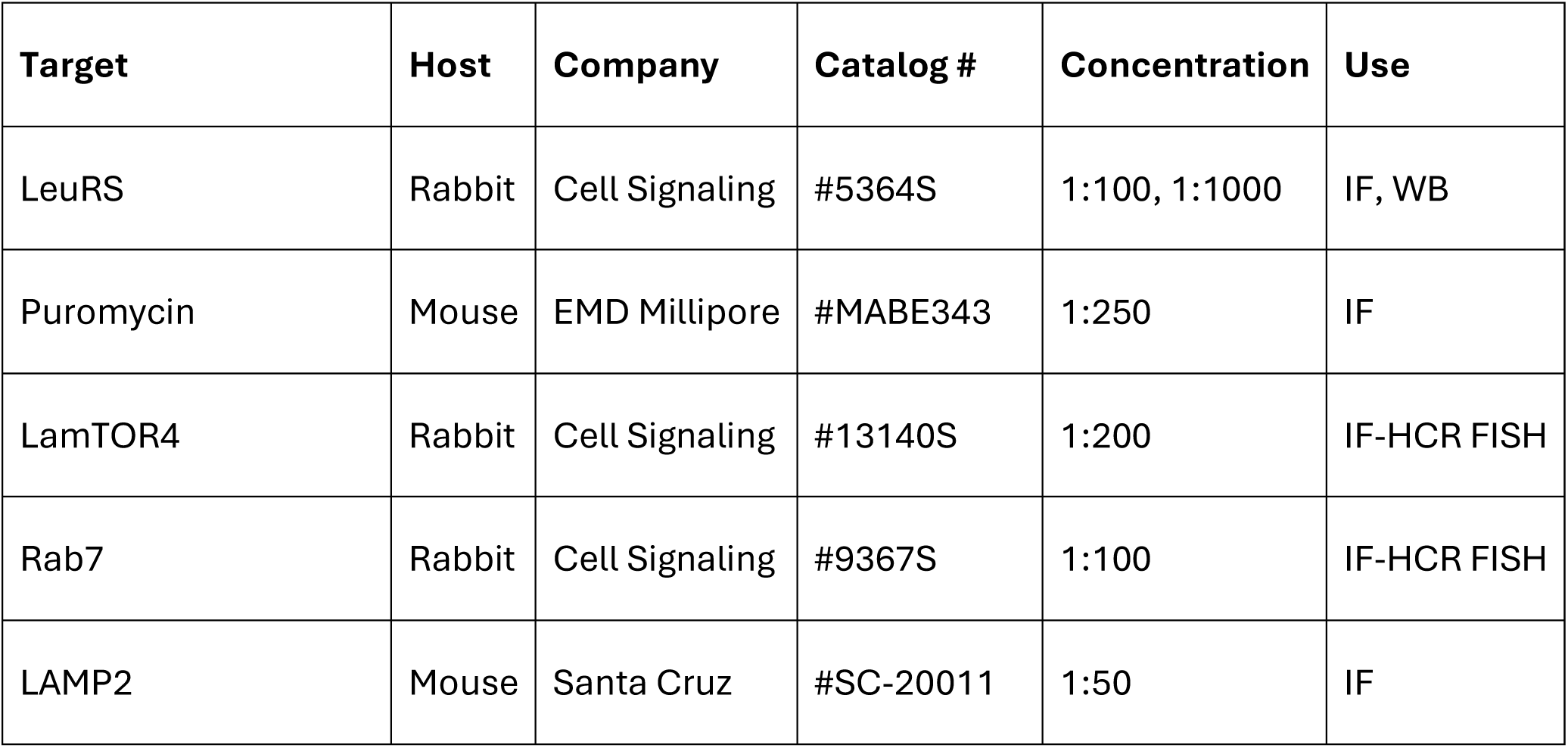
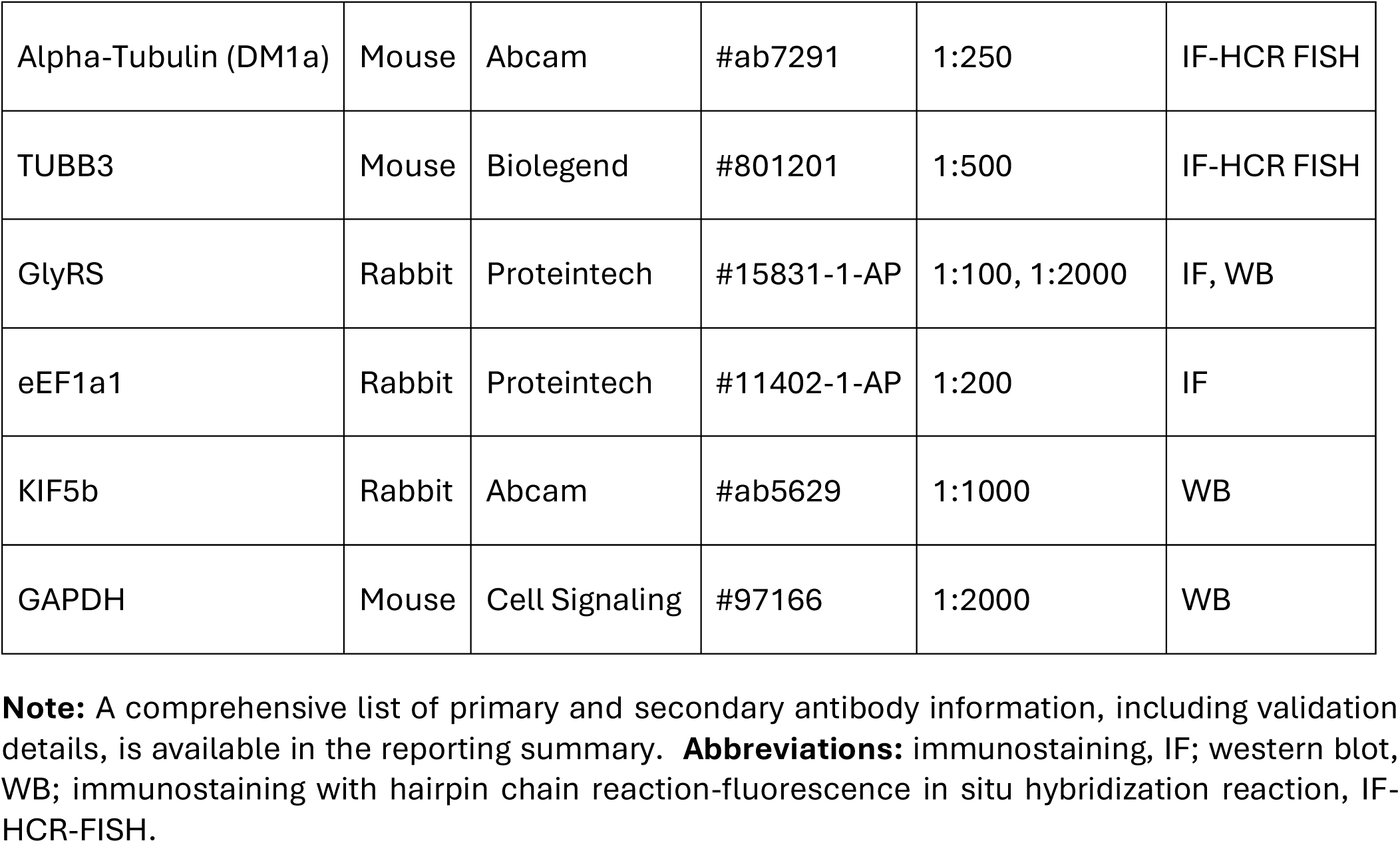

## Data and Code Availability

All data supporting the findings of this study are available within the paper, its Supplementary Data, and associated source data files. Custom code used for data analysis with standard software packages (ImageJ, MATLAB) and additional raw and processed data generated are available from the corresponding author upon reasonable request.

## Acknowledgments

Funding for this work was provided by the National Institutes of Health (NIH) R01s-HL133080 and HL149891 to BLP., F32HL170583 and T32AR053461 to JMP. BLP also awarded Israel Binational Science Foundation (BSF) Award 2019126 and the Foundation Leducq Research grant no. 20CVD01. KU was funded by the American Heart Association Career Development Award (24CDA1267507).

## Contributions

JMP and BLP conceived the project. JMP wrote the manuscript and performed the majority of experiments. BLP aided experimental design and data review and interpretation. BC aided in the generation of labeled tRNAs for live cell experiments. JMP and KU performed washout experiment analysis and KU developed MATLAB pipeline for RNA distribution analysis and recovery index. JMP and VC performed Weka analysis of tRNA microtubule colocalization and imaging. CB and ML performed STORM imaging and analysis. AB performed neuronal isolations and neuronal western blots.

