## Supplemental Data Figures and Legends for "Active transport of tRNAs facilitates distributed protein synthesis"

**Running Title**: Active transport of tRNAs facilitates distributed protein synthesis

**Supplementary Data and Legends**

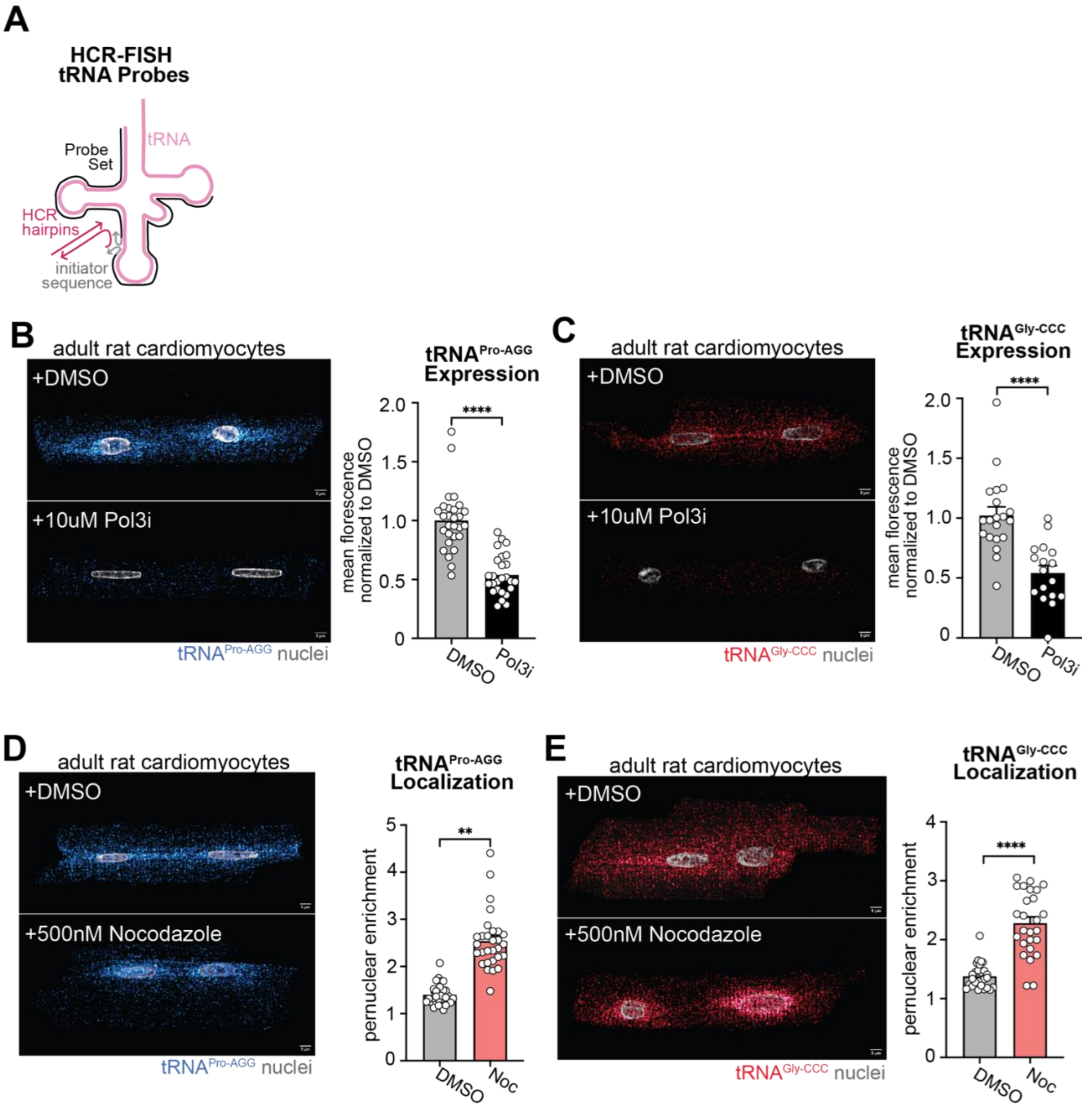


**Figure S1: Validation of additional single tRNA probes.** (A) Schematic of tRNA probe design for a single tRNA. (B) $tRNA^{Pro\text{-}AGG}$ (blue, $N$ = 3) and (C) $tRNA^{Gly\text{-}CCC}$ ($N$ = 2) representative images and quantification of mean fluorescence intensity (MFI) normalized to DMSO in ARVMs treated for 96h with DMSO (grey bar) or a Polymerase III inhibitor (10 μM Pol3i, black bar). (D) $tRNA^{Pro\text{-}AGG}$ (blue) and

(E) $tRNA^{Gly\text{-}CCC}$ representative images and quantification of perinuclear enrichment of mean fluorescence intensity relative to cytoplasmic levels in ARVMs treated for overnight with DMSO (grey bar) or a 500nM nocodazole (Noc, light red bar).All experiments were done in biological triplicate (*N* = 3), with 10 cells imaged per biological replicate (*n* = 10), unless noted. Data are presented as the mean ± SEM. *P < 0.05, **P < 0.01, ***P < 0.001, ****P < 0.0001. Two-sided Student's t-test.

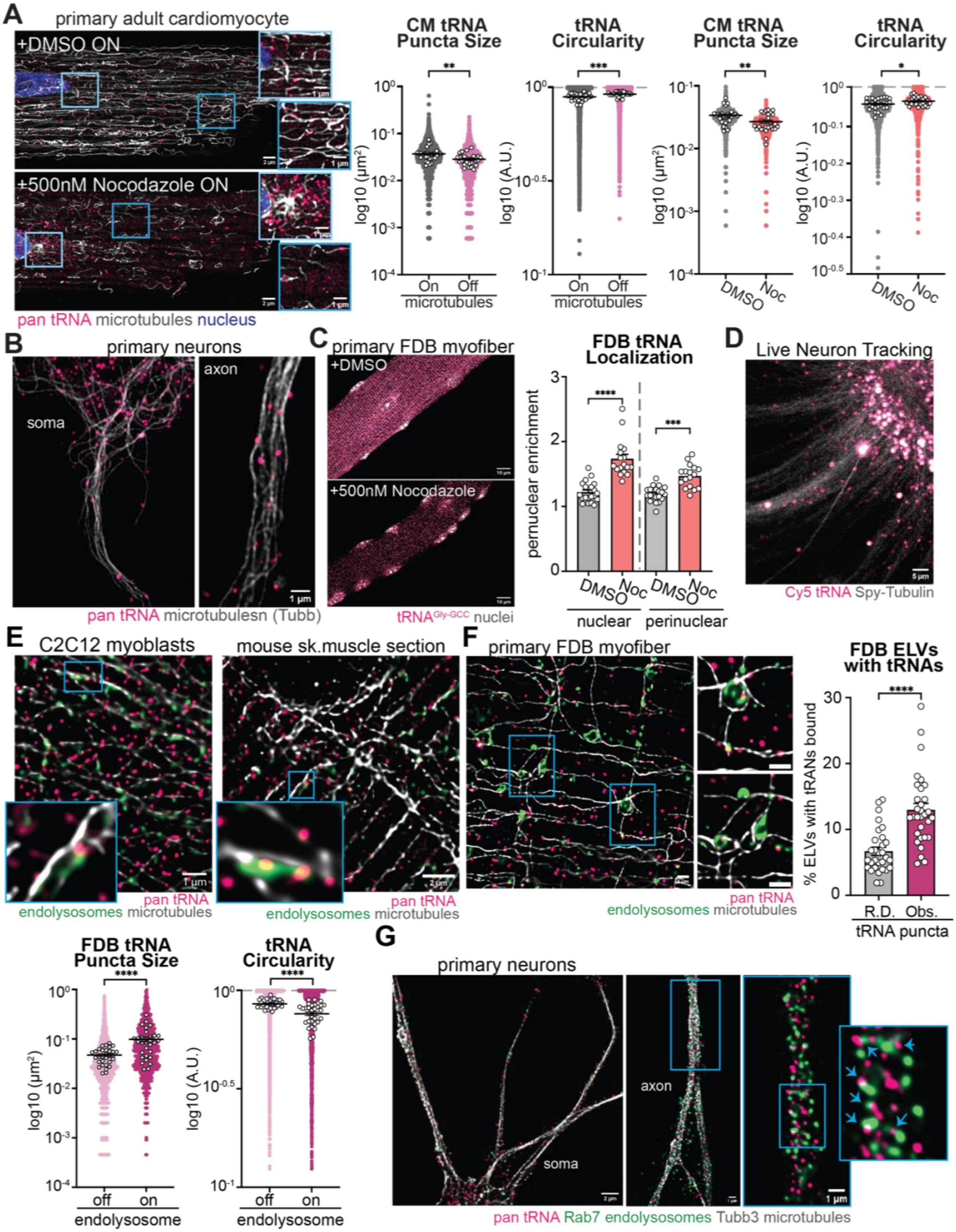
A
primary adult cardiomyocyte
+DMSO ON
+500nM Nocodazole ON
pan tRNA microtubules nucleus
CM tRNA Puncta Size
log10 (µm²)
On Off
microtubules
tRNA Circularity
log10 (A.U.)
CM tRNA Puncta Size
DMSO Noc
tRNA Circularity
B
primary neurons
soma
axon
pan tRNA microtubulesn (Tubb)
C
primary FDB myofiber
+DMSO
+500nM Nocodazole
tRNAGly-GCC nuclei
FDB tRNA Localization
pernuclear enrichment
nuclear
perinuclear
D
Live Neuron Tracking
Cy5 tRNA Spy-Tubulin
E
C2C12 myoblasts
mouse sk.muscle section
pan tRNA
endolysosomes microtubules
F
primary FDB myofiber
FDB ELVs with tRNAs
% ELVs with tRANs bound
R.D. Obs.
tRNA puncta
FDB tRNA Puncta Size
off on
endolysosome
tRNA Circularity
G
primary neurons
axon
soma
pan tRNA Rab7 endolysosomes Tubb3 microtubules

**Figure S2: tRNAs undergo microtubule-mediated transport on the outside of endolysosomes in adult skeletal muscle myofibers, cardiomyocytes, and neurons.** (A) Representative images of tRNA (pink) and DM1a-stained microtubules (silver) in ARVMs treated overnight with DMSO or 500nM nocodazole (Noc). Blue box insets indicate areas of zoom and quantifications of pan tRNA puncta size and circularity (plotted on the log10 scale) when located on microtubules (grey), off microtubules (pink), following overnight DMSO (light grey) or nocodazole (red) treatment in ARVMs. Each individual graphical dot indicates an individual puncta measurement, and white dots indicate the means for each individual cell. (B) Representative images of endogenous Tubb3 + microtubules (grey) and pan tRNAs (pink) in primary, postnatal, cortical neuron soma and axon. (C) Representative images of endogenous tRNAs (pink) in FDB myofibers treated with DMSO or 500nM nocodazole overnight and quantification of nuclear and perinuclear tRNA enrichment relative to cytoplasmic levels. (D) Snapshot of live-cell movie in a primary postnatal cortical neuron where the microtubules are stained in SpyTubulin (silver), and cells have been transfected with Cy5-tRNAs (pink). (E) Representative images of C2C12 myoblasts, a mouse T.A. skeletal muscle section and (F) isolated primary rat FDB myofiber with staining for DM1a labeled microtubules (silver), pan tRNAs (pink), and LamTOR labeled ELVs (green) with FDB quantifications of ELVs with tRNAs bound (dark pink, Obs.) compared to randomized puncta of the same size and shape (Randomized, R.D., grey). From FDB tRNA punctum measurements, quantifications of area and circularity of tRNAs on (dark pink) and off (light pink) the ELVs. Data are plotted on the log10 scale with each individual graphical dot indicates an individual puncta measurement, and white dots indicate the means for each individual cell. (G) Representative images of endogenous pan tRNAs (pink), Tubb3-stained microtubules (silver), and Rab7-stained ELVs (green) in primary neurons. Blue arrows indicate tRNA-ELV colocalization. All experiments were done in biological triplicate (*N* = 3), with 10 cells imaged per biological replicate (*n* = 10), unless noted. Data are presented as the mean ± SEM. *P < 0.05, **P < 0.01, ***P < 0.001, ****P < 0.0001 as determined with Two-sided Student's t-test.

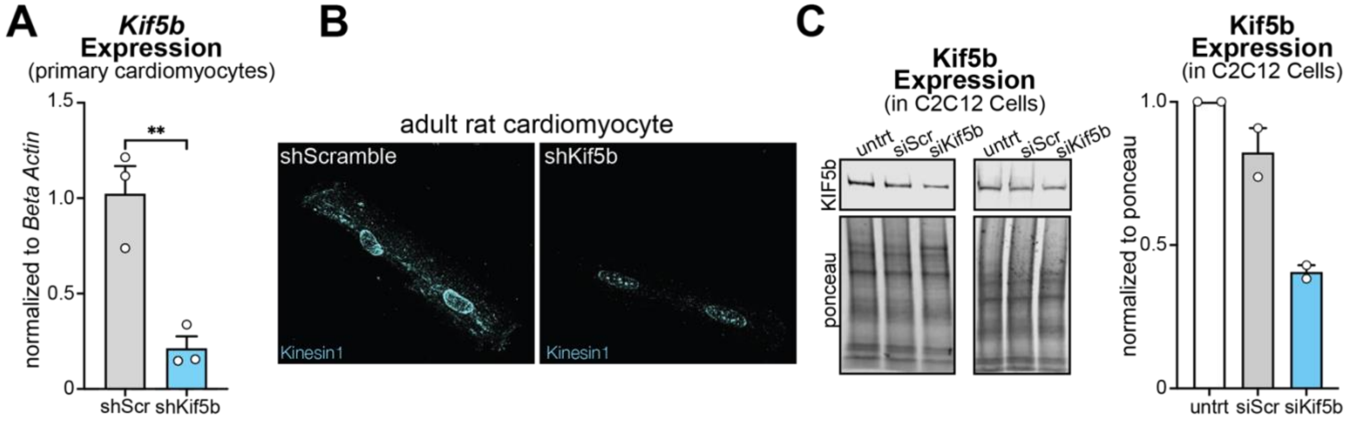


**Figure S3: Knockdown of Kinesin 1.** (A) *Kif5b* mRNA expression, relative to beta actin loading control, from three biological preps of adult rat cardiomyocytes in cells receiving an adenovirus targeting a scramble sequence (grey) or targeting Kif5b (light blue) and (B) KIF5b staining in those cells. (C) Kif5b protein expression, normalized to total protein by ponceau stain) and quantification from two biological preps of C2C12 cells following 50 nM siRNAs targeting a scramble sequence (siScr, grey), Kif5b (siKif5b, light blue), or untransfected controls (untrt, white) after 72h. Data are presented as the mean ± SEM. $^*P < 0.05$, $^{**}P < 0.01$, $^{***}P < 0.001$, $^{****}P < 0.0001$ as determined by Two-sided Student’s t-test.

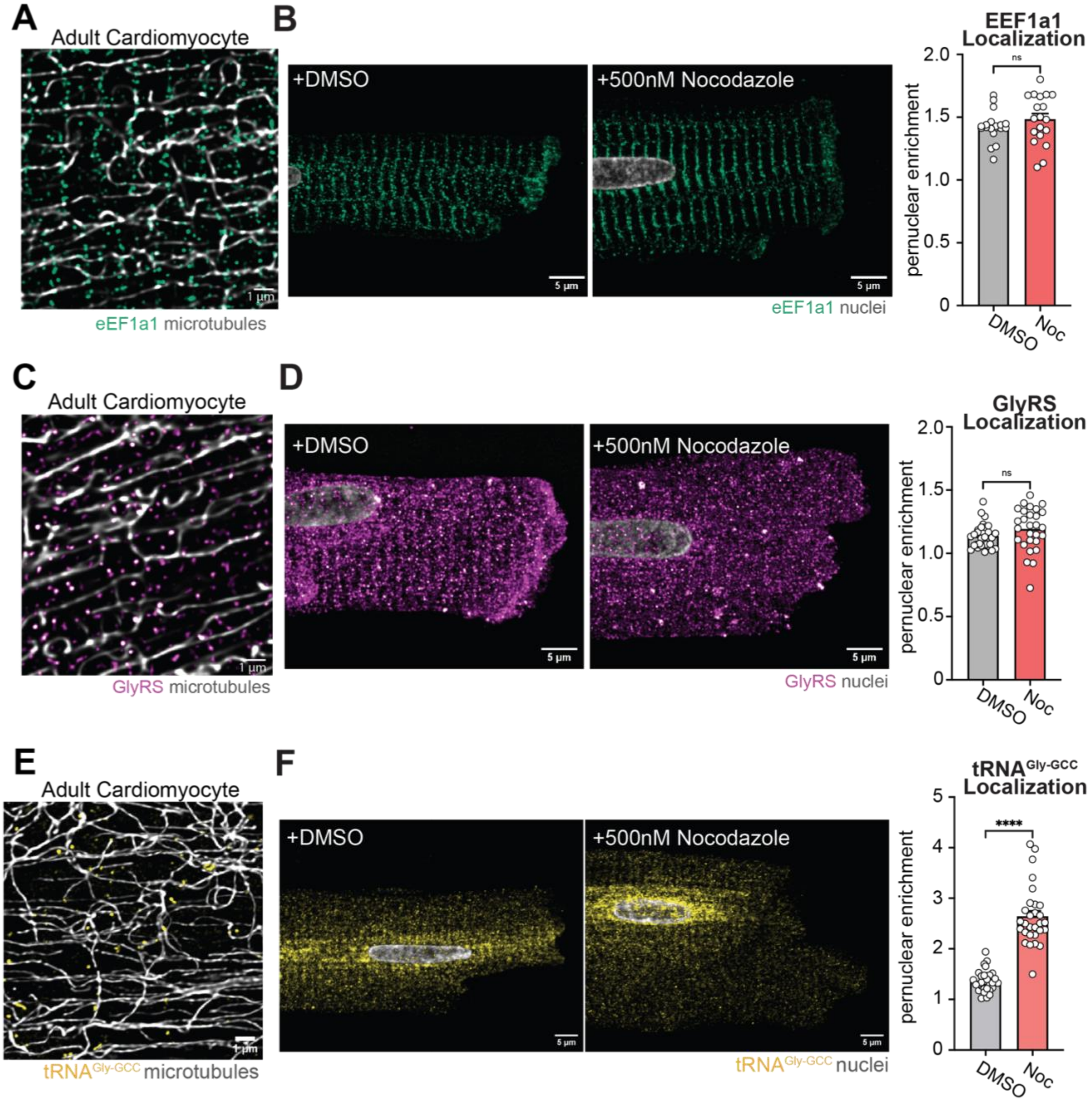


**Figure S4: The tRNA binding proteins EEF1a1 and GlyRS are not nocodazole sensitive.** (A) Representative images of eEF1a1 (green) and DM1a-stained microtubules (silver) in ARVMs. (B) Representative images of eEF1a1 (green) in ARVMs treated overnight with DMSO (grey) or 500nM nocodazole (light red) and quantification of perinuclear mean fluorescence intensity relative to cytoplasmic levels. (C) Representative images of GLyRS (purple) and DM1a-stained microtubules (silver) in ARVMs. (D) Representative images of GlyRS (purple) in ARVMs treated overnight with DMSO (grey) or 500nM nocodazole (light red) and quantification of perinuclear mean fluorescence intensity relative to cytoplasmic levels. (E) Representative images of $tRNA^{Gly\text{-}GCC}$ (yellow) and DM1astained microtubules (silver) in ARVMs. (F) Representative images of $tRNA^{Gly\text{-}GCC}$ (yellow) in ARVMs treated overnight with DMSO (grey) or 500nM nocodazole (light red) and quantification of perinuclear mean fluorescence intensity relative to cytoplasmic levels. All experiments were done in biological triplicate (*N* = 3), excluding b, done in duplicate (*N* = 2), with 10 cells imaged per biological replicate (*n* = 10), unless noted. Data are presented as the mean ± SEM. **P < 0.05, **P < 0.01, ***P < 0.001, ****P < 0.0001. Two-sided Student's t-test.

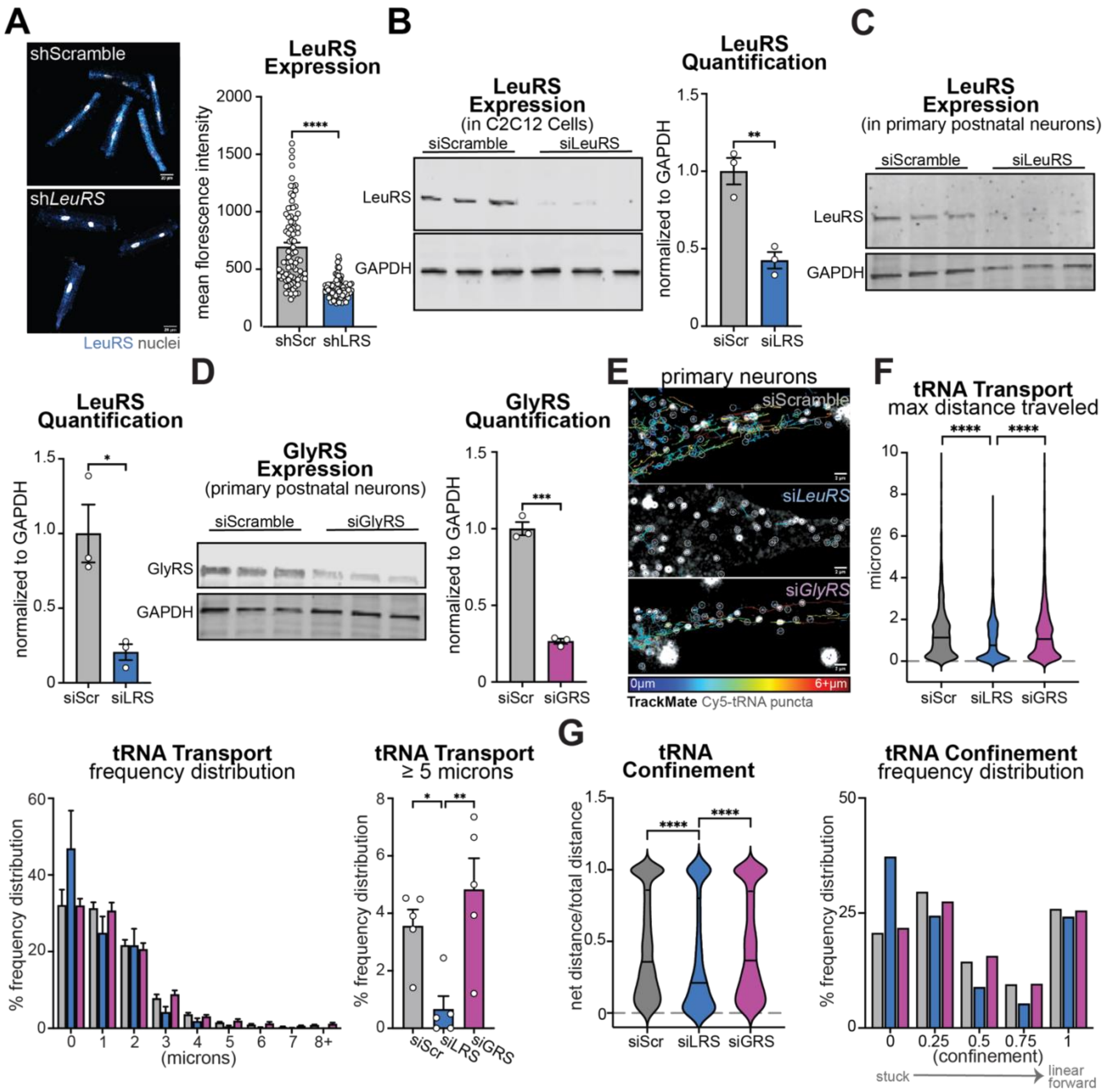


**Figure S5: Determining adaptors for long-range tRNA transport.** (A) Representative images of ARVMs stained for LeuRS (blue staining) following 96h of treatment with an adenovirus encoding short hairpins targeting a scrambled sequence (shScramble, shScr, grey bar) or targeting the knockdown of Leucyl-tRNA synthetase (*LeuRS*, sh*LeuRS*, shLRS, blue bar) and quantification of LeuRS mean florescence intensity. (B) Western blot in C2C12's treated for 96h with 50nM siRNAs to knockdown LeuRS (siLRS, blue) or a scramble sequence (siScr, grey) and quantification of LeuRS expression normalized to GAPDH. (C) Western blot in primary adult neurons of LeuRS expression following 96h post-delivery of 50nM siRNAs to knockdown LeuRS (siLRS, blue) or a scramble sequence (siScr, grey) and quantification of LeuRS expression. (D) Western blot in primary adult neurons of Glycyl-tRNA synthetase (GlyRS) expression following 96h post-delivery of 50nM siRNAs to knockdown GlyRS (siGRS, purple) or a scramble sequence (siScr, grey) and quantification of GlyRS expression. (E) Representative TrackMate tRNA movements in primary neurons following 96h of transient transfection with 50nM siScramble (siScr, grey), siLeuRS (siLRS, blue), or siGlyRS (siGRS) and delivery of Cy5-tRNAs. (F) Quantification of maximum distance traveled per tRNA punctum, associated frequency distribution plots, and percentage of events greater than 5 microns. (G) Quantification of confinement ratio (where 0 is confined, and 1 is linear movements) of tRNA punctum and associated frequency distribution plots. LeuRS ARVM and C2C12 experiments were performed in biological triplicate, analyzing more than 50 cells/biological replicate. Neuronal experiments are done in biological duplicate preps, analyzing over 100 puncta/prep. Data are presented as the mean ± SEM. *P < 0.05, **P < 0.01, ***P < 0.001, ****P < 0.0001 as determined by Two-sided Student's t-test (a, c-g) or one-way ANOVA with post hoc Tukey's multiple comparisons (h-i).

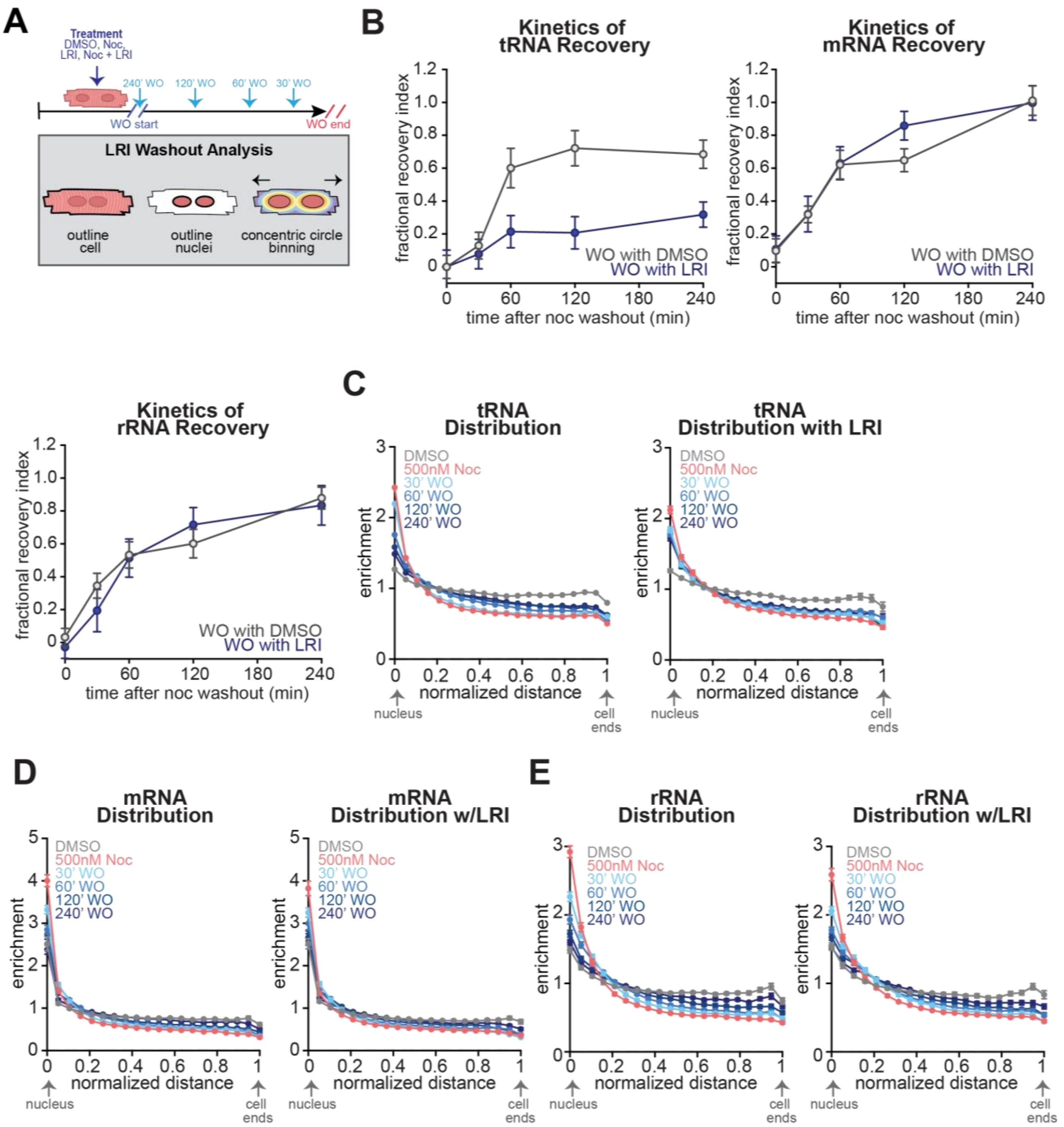


**Figure S6: Blocking LeuRS-endolysosome interactions specifically disrupts tRNA transport**. (A) Schematic of LeuRS-RagD inhibitor nocodazole washout experiments and concentric circle binning analysis used. (B) tRNA, mRNA, and 28s rRNA quantifications of fractional recovery index (50% RNA fractional recovery) of each respective RNA with (blue line, LRI) or without (grey line, DMSO) the LRI in washout media. (C-E) Quantifications of tRNA, mRNA, and rRNA, with or without LRI in washout, cumulative frequency distribution plots of each RNA's redistribution from the outside of the nucleus to the edges of the cell. All experiments were done in biological triplicate ($N$ = 3), with 10 cells imaged per biological replicate ($n$ = 10). Data are presented as the mean ± SEM.

**Supplementary Video Legends**

**Movie S1: Cy5-tRNA (pink) trafficking in C2C12 myotubes.** Cy5-tRNAs (pink) transiently transfected into C2C12 myotubes and live cell imaged using Airyscan microscopy.

**Movie S2: tRNAs that can and cannot be charged undergo long-range transport.** Cy3-tRNAs (from yeast, pink) and Cy5-tRNAs (from e.coli, blue) trafficking in C2C12 myotube.

**Movie S3: tRNAs traffic on microtubules in myotubes.** Cy5-tRNAs (pink) traffic on SPY-tubulin stained microtubules (silver) in C2C12 muscle cells.

**Movie S4: tRNAs traffic on microtubules in neurons.** Cy5-tRNAs (pink) traffic on SPY-tubulin stained microtubules (silver) in primary neurons.

**Movie S5: Microtubule disruption by nocodazole treatment impairs tRNA movement.** Cy5-tRNAs (pink) transiently transfected into C2C12 myotubes labeled with the live cell microtubule dye SPY-tubulin (grey) overnight. The cell in image (left) was first treated with DMSO, and then treated with (right) 500 nM nocodazole for 30 minutes. The microtubule network begins to break down, as demonstrated by the loss of SpyTubulin signal, and tRNA trafficking is halted.

**Movie S6. tRNAs hitchhike on endolysosomes in muscle.** Cy5-tRNAs (pink) transiently transfected into C2C12 myotubes labeled with 50nM endolysosome dye LysoTracker (Green) for 1 hour. DMSO treatment (left) or 500nM nocodazole treatment (right).

**Movie S7: tRNAs hitchhike on endolysosomes in neurons.** Cy5-tRNAs (pink) hitchhike on Lysotracker-stained endolysosomes (green) in primary neurons.

**Movie S8: Electroporated Cy5-tRNAs hitchhike on endolysosomes in C2C12 cells.** Cy5-tRNAs (pink) electroporated into C2C12 myotubes and labeled with the live cell endolysosome dye LysoTracker (Green) for 1 hour.

**Movie S9: Kinesin-1 depletion reduces long-range tRNA transport.** Cy5-tRNAs (pink) and Lysotracker-stained endolysosomes (green) in C2C12 cells treated for 72h with 50 nM scramble siRNAs (left) or siRNAs targeting *Kif5b* (right).

**Movie S10: Loss of LeuRS reduces long-range tRNA transport.** Cy5-tRNAs (pink) and Lysotracker-stained endolysosomes (green) in C2C12 cells treated for 96h with 50 nM scramble (left) or *LeuRS* targeting siRNAs.

**Movie S11: Loss of LeuRS, but not GlyRS reduces long-range tRNA transport primary in neurons.** Cy5-tRNAs (pink) transfected into primary neurons treated for 6 days with 50 nM siRNAs encoding a scramble sequence (top), or targeting *LeuRS* (middle) and *GlyRS* (bottom).
